# Metalloprotease-driven remodeling of PTK7 and the cell surfaceome promotes metastatic fitness of circulating colorectal tumor cells

**DOI:** 10.64898/2026.07.24.740526

**Authors:** Olivier Cabaud, Anaïs Aulas, Alexia M. Lopresti, Claire Acquaviva, Pascal Finetti, Charlotte Dessaux, Laetitia Ganier, Quentin DaCosta, Claire Germier, Lenaïg Mescam, Abdessamad Elkaoutari, Stéphane Audebert, Luc Camoin, Bernadette de Rauglaudre, Laurys Boudin, Emilie Denicolai, Gwenaël Lumet, Agathe Cohendet, Maëlle Picard, David Birnbaum, Caroline Gouarne, Brice Chanez, Cécile de Chaisemartin, Bernard Lelong, Sylvie Marchetto, Anthony Gonçalves, Daniel Birnbaum, François Bertucci, Jean-Paul Borg, Emilie Mamessier

## Abstract

Circulating tumor cells (CTCs) are the potential seeds of distant metastases; however, little is known about how they survive in the bloodstream. Using a large cohort of colorectal cancer (CRC) patients, we found that the pseudokinase receptor PTK7 is highly expressed in primary tumors and metastatic lesions. Consistent with previous reports, high PTK7 expression is associated with reduced disease-free survival and increased metastatic dissemination. Surprisingly, PTK7 is absent from most CTCs and undergoes a cell-autonomous ON^tumor^/OFF^CTC^/ON^metastasis^ switch that can be recapitulated in a xenografted mouse model, in *in vitro* systems, and a fluidic platform. PTK7-negative cancer cells exhibit increased expression of YAP1-driven genes, senescence-like features, and enhanced resistance to hemodynamic stress following loss of cell-cell and cell-matrix adhesion. This adaptive phenotype depends on metalloproteases, notably ADAM17, whose cleavage activity remodels the CTCs surfaceome. Functionally, the PTK7 OFF^CTC^ state confers enhanced metastatic potential *in vivo*, and can be pharmacologically suppressed using metalloprotease inhibitors. Collectively, our findings identify a reversible, cell-autonomous, protease-driven surfaceome remodeling program that enables metastatic adaptation during hematogenous dissemination.

**Highlights / statement of significance:** By investigating potential markers for circulating colorectal tumor cells with strong metastatic potential, we describe a reversible and cell-autonomous remodeling of the circulating tumor cell surfaceome in patients that confers resistance to anoikis and stress induced by entry into the bloodstream.

**One Sentence Summary:** The dynamic regulation of PTK7 serves as a surrogate marker for tumor cell plasticity, aggressiveness, survival in the bloodstream, and efficiency in forming metastases.

**Trial registration:** CTC colon Cohort: registered on https://ClinicalTrials.gov identifier NCT03256084; date of registration 2017-07-17

B-Org cohort: registered on https://ClinicalTrials.gov NCT05384184; date of registration 2019-06-06

**Ethics statement for animal experiments:** Studies on animals were conducted in accordance with the current ethical standards of the European Community (Directive 2010/63/EU), the Ethics Committee for Animal Experimentation (CEEA#14) and the French Ministry of Higher Education and Research, which approved and authorized the entire procedure described in this paper (project number APAFIS #35294).

## INTRODUCTION

Colorectal cancer (CRC) is highly metastatic, and its high mortality rate is due to metastases affecting distant organs (1). The occurrence of metastasis could notably be anticipated using the Consensus Molecular Subtype (CMS) classification, which categorizes primary CRC into four subtypes with different metastatic risks (2); however, this classification has not yet been implemented in clinical practice. Biomarkers that can predict and prevent early metastasis development are therefore still urgently needed.

Circulating tumor cells (CTCs) are tumor cells that detach from the original tumor and enter the bloodstream. They are considered as potential seeds for metastasis (3). Because epithelial cells are not adapted to flow conditions, the vast majority of CTCs do not survive the stressful conditions in the bloodstream and do not reach distant sites to form metastatic foci (4–6). Quantity, *i.e.* a high number of CTCs released into the blood, and quality, *i.e.* CTCs with increased survivability, are factors that may favor the occurrence of metastases (7). While the quantity factor is well documented in cancers, the quality factor is less understood due to the difficulty in identifying and characterizing CTCs among blood cells, and in properly assessing their heterogeneity (8,9). Enumeration provides only part of the relevant information about CTCs, which probably explains the low clinical added value of CTC counts compared to other clinicopathological variables.

Not all tumor cells detected in the bloodstream have metastatic potential (10). A better understanding of the biological characteristics of CTCs as they progress toward their niche could enable prediction, and potentially allow targeting and limiting the development of metastases. The cytological description of CTCs isolated from patients with CRC, either as clusters or single cells, and the expression of molecular markers related to tumor aggressiveness, such as EGFR, LGR5, and CD44v6, have thus been investigated (9,11–13). Some of these markers may be useful for tailoring therapies in metastatic CRC (mCRC) (9,14). However, very few studies have investigated markers expressed by CTCs in primary CRC patients that correlate with the development of metastases (15,16).

PTK7 is a catalytically inactive member of the receptor tyrosine kinases that is highly expressed in CRC cells (17). Its overexpression in ≈35% of primary CRCs correlates with increased incidence of metastases and reduced mouse and patient survival (18). *In vitro*, PTK7 expression in tumor cells is associated with stem cell potential (19), and enhanced migratory capacity (18,20,21). Together, these observations suggest that PTK7 is a promising target for therapeutic intervention in CRC (22–24).

In this study, we investigated the contribution of PTK7 to metastasis development in CRC, and revealed a dynamic and transient “switch-like” regulation of PTK7 expression in tumor cells that is critical for the survival of CTCs in the blood. This regulation is primarily mediated by metalloprotease activity that remodels the tumor cell surfaceome, leading to PTK7 cleavage, as well as YAP1-dependent and senescence-like features. Altogether, our results shed light on a phenotype associated with CTC plasticity and its importance during the metastatic cascade.

## RESULTS

### PTK7 is upregulated in primary and metastatic colorectal cancer

To explore PTK7 expression in primary and metastatic CRCs on a larger scale than previous studies that reported contradictory results (18,25), we created a database of clinically annotated gene expression profiles (**Tables S1**, **S2**) that included 95 normal colon (NC) samples, 2,239 primary CRC samples, and 68 metastatic CRC (mCRC) samples. *PTK7* mRNA levels were heterogeneous in the 2,239 primary tumors (∼8 units on the log_2_ scale) and in the metastatic samples (∼6 units on the log_2_ scale). Compared to NC, *PTK7* mRNA was upregulated in primary CRC (p=1.06E-34) and mCRC (p=8.5E-28), and *PTK7* was also upregulated in mCRC compared to primary CRC (p=3.63E-04) (**Figure 1A**).

**Figure 1:**
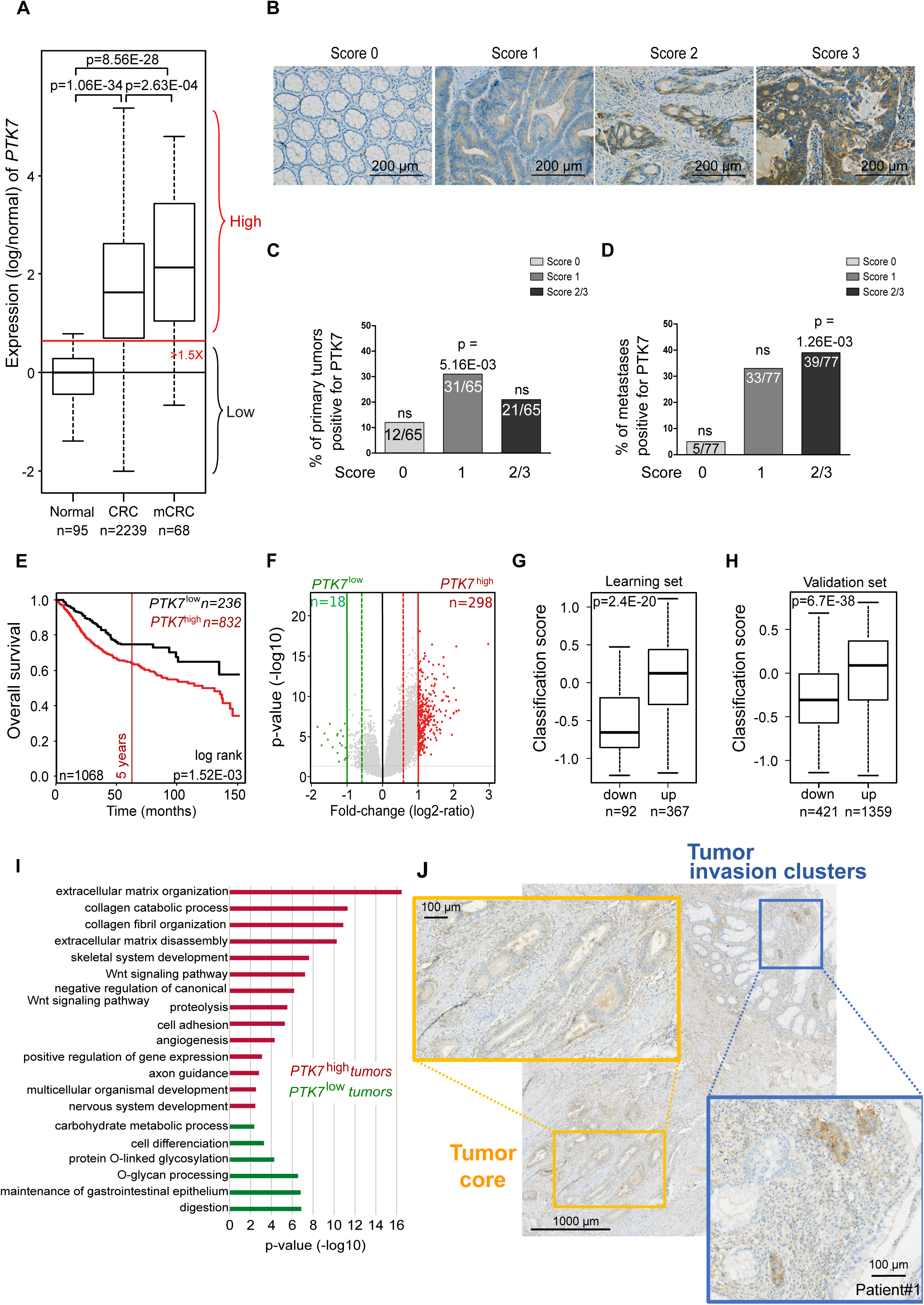
Overexpression of PTK7 in CRC is an independent poor overall survival marker for patients with CTC and is associated with extracellular matrix remodeling. **A.** Boxplots showing *PTK7* mRNA levels (log^2^) in NC samples (n = 95), in CRC primary tumors (CRC; n = 2,239) and in CRC metastases (mCRC; n = 68). Expression was normalized to NC samples. PTK7^High^ samples had expression above 1.5 (horizontal red line) and *PTK7^Low^* samples correspond to samples with expression below 1.5. Median and ranges are indicated for each boxplot. *PTK7* expression was compared between groups using ANOVA (multiple comparisons). **B.** Immunohistochemistry (IHC) for PTK7 in an adjacent NC tissue (Score 0) and three primary CRC tumors (Score 1 to Score 3) FFPE sections**. C-D.** Scoring of PTK7 staining in primary CRC tumor samples (C) or CRC metastases (D). A binomial exact test was used to evaluate the enrichment in each score (one side alternative hypothesis: enrichment higher than 1/3). **E.** Kaplan-Meier curves for OS of patients with *PTK7^High^* (red curve) or and *PTK7^Low^* (black curve) primary CRC. The p-value is for the log-rank test. **F.** Volcano plot showing the 316 genes differentially expressed between the *PTK7^High^ and PTK7^Low^* tumors in the learning set (TCGA, n = 459 samples). Genes upregulated and those downregulated in the *PTK7^High^* samples are colored red and green, respectively (p<5%, q<25% & |FC| > 2x). **G-H.** Metagene-based prediction score in *PTK7^High^* samples compared to those of PTK7^Low^ samples in the learning set (G) and in the independent validation set (H). The p-values are for the Student t-test. I. Gene ontologies associated with *PTK7^High^* tumors (red bars) or *PTK7^Low^* tumors (green bars). The p-values (-log10) were corrected for multiple testing using the Benjamini-Hochberg method**. J.** Expression of PTK7 by IHC in tumor cells located in the tumor core (yellow square) or in tumor invasive clusters (blue square).

To assess whether PTK7 protein expression reflected the *PTK7* mRNA trend, we performed immunohistochemistry (IHC) (**Figure 1B**) on formaldehyde-fixed paraffin-embedded (FFPE) specimens from 65 patients with primary CRC and 77 patients with metastatic disease (CTC-colon and B-Org cohorts). PTK7 was expressed in 81% of primary tumors (53 of 65), with high expression (intensity score 2 or 3) observed in 32% (21 of 65) of samples (**Figure 1B-C**). Adjacent NC tissues, when present, were negative for PTK7 (**Figure 1B**), except at the base of crypts (**Figure S1A**) (19). PTK7 was also detected in 93% of metastatic specimens (72 of 77), with 51% (39 of 77) showing high expression (score 2/3) (**Figure 1D**). Most primary CRC samples had a score of 1, whereas most mCRC samples had scores of 2 or 3 (**Figure 1C-D**).

Thus, most CRC cases, especially metastases, show higher PTK7 expression than NC samples. This is the first evidence that PTK7 is strongly expressed at both the mRNA and protein levels in mCRC.

### Overexpression of *PTK7* in CRC is an independent factor for poor overall survival

We categorized CRC tumors in our dataset as *PTK7^High^*(n=1,501), if the ratio of *PTK7* mRNA level in the tumor to the *PTK7* mRNA median in the 95 NC samples was >1.5, or as *PTK7^Low^* (n = 429), if the ratio was <1.5. We then correlated the *PTK7^High^* or *PTK7^Low^* classifications with clinicopathological and molecular characteristics of the 2,239 CRC samples. We did not find any correlation with patient age, tumor location, tumor stage, tumor grade, or tumor mismatch repair (MMR) status, but we found correlations with patient gender (p = 0.0482) and CMS classification (p = 1.52 E-42) (**Table S2**) (2). Compared to *PTK7^Low^*, *PTK7^High^* tumors more frequently belong to the CMS2 WNT-activated class (canonical), and CMS4 TGF-β-activated class (mesenchymal), (ANOVA test, p=5.6E-89, **Figure S1B**).

Next, we investigated whether high *PTK7* mRNA levels were associated with the prognosis of CRC patients undergoing surgery for their primary tumor. The overall survival (OS) data were available for 1,068 of the 2,239 CRC cases. Among these 1,068 cases, the median follow-up was 39 months (range 1-212) and the 5-year OS rate was 68% (CI 64-71; **Table S2**). *PTK7^Low^* cases had a better 5-year OS rate than *PTK7^High^* cases (75% *vs* 66%, p=1.52 E-03, **Figure 1E**). Considering the variability of the *PTK7^High/^PTK7^Low^* distribution among CMS classes, we analyzed the OS of each CMS class with regard to *PTK7* expression using the log-rank test (p=1.44 E-05). Among patients with CMS4 CRC (*i.e.,* tumors with mesenchymal features and higher metastatic risk), the OS of those with *PTK7^High^* tumors was 16% lower than those with *PTK7^Low^* tumors (p= 0.06, **Figure S1C**).

Univariate prognostic analysis showed a hazard ratio (HR) for OS event of 1.53 (95% CI 1.15-2.04) in the *PTK7^High^* class compared to the *PTK7^Low^* class (p=3.76E-03, Wald test) (**Table S3**). Other significant variables were pathological stage (p=1.35E-51), grade (p= 1.42E-03), and CMS status (p= 4.53E-08). Patient age, gender, tumor location, and MMR status were not associated with OS. In multivariate analysis, *PTK7^High^/PTK7^Low^*classification remained associated with OS (p=1.47E-02, Wald test), as did pathological stage and CMS classification (**Table S3**).

Our data show that high expression of *PTK7* in CRC is an independent poor-prognostic factor for OS in CRC patients, especially those of the CMS4 class, who are at higher risk of developing metastases.

### Tumors that overexpress PTK7 exhibit pro-invasive mesenchymal features rather than epithelial features

To study the biological functions associated with *PTK7^High^*expression in primary CRC, we compared the transcriptomic profiles of 92 *PTK7^Low^*and 367 *PTK7^High^* tumors in the TCGA dataset (learning set) and identified 316 differentially expressed genes (DEGs), including 298 overexpressed and 18 underexpressed genes in *PTK7^High^* tumors (**Figure 1F**; **Table S4**). A metagene was derived from these 316 DEGs (see Methods section) and used to classify the samples. As expected, this metagene efficiently classified the 459 samples in the learning set (p = 2.48E-20, t-test; **Figure 1G**). Importantly, its robustness was confirmed in an independent validation group of 1,780 samples (p = 6.70E-38, t-test; **Figure 1H**).

Ontology analysis of the 316 DEG revealed increased expression of genes involved in extracellular matrix (ECM) reorganization (p = 3.45E-17), collagen degradation (p = 5.17E-12), negative regulation of canonical WNT signaling (p = 6.51E-07), cell adhesion (p = 5.11E-06), angiogenesis (p = 4.80E-05), and axon guidance (p = 3.00E-04) in *PTK7^High^* tumors (**Figure 1I**, **Table S5**). Specifically, compared to *PTK7^Low^* tumors, *PTK7^High^*tumors were enriched for several metagenes associated with EMT and tumor cell budding (GSEA EMT hallmark metagene V62, EMT/CHL model, EMT score, tumor cell budding signature, and C1 to C4 EMT states) (**Figure S1D**). Overall, the major biological pathway activated in *PTK7^High^* tumors was related to cell-substrate interactions through ECM remodeling, a hallmark of cancer cell invasion, migration, and metastasis. Moreover, PTK7 was highly expressed in tumor invasion clusters in CRC (**Figure 1J, S1E**). In addition, PTK7 expression was up-regulated in HCT116 or SW480 cell lines exposed to epithelial-mesenchymal transition (EMT)-inducing agent and in migrating cells submitted to a scratch assay (**Figure S1F-G**). These data are consistent with previous reports of PTK7 involvement in tumor cell migration and confirm expression of PTK7 as a marker of the early migratory steps of the metastasis cascade within the tumor (18,20,26).

We therefore decided to evaluate PTK7 expression in the subsequent step of metastatic dissemination, *i.e.* in CTCs.

### In contrast to primary and metastatic CRC cells, PTK7 is rarely found on CTCs

To evaluate PTK7 expression in CRC-derived cells circulating in the bloodstream, we isolated CTCs from CRC patients (**Table S6**) based on their size and deformability, independently of any specific cell markers. After isolation, CTCs were identified by their morphology, size, and phenotype, determined by spectral immunofluorescence (IF) (*i.e.,* CD45-negative and CK^+^ and/or EPCAM^+^ and/or VIM^+^ cells) (**Figure 2A** and **S2A**). We analyzed blood samples from 38 CRC patients whose primary tumors were positive for PTK7 by IHC (score 1 to 3). We enumerated CTCs (mean: 31 CTCs/mL; range: 0.3 - 251 CTCs/mL) and determined the percentage of CTCs expressing PTK7. PTK7-positive (PTK7^+^) CTCs were detected in 7 out of 38 patients (18.4%), but not all CTCs in these seven samples were positive for PTK7 (range: 0.5% to 30%) (**Figure 2B**, CRC). Among the 6,737 CTCs identified, only 47 were PTK7^+^ and 6,690 were PTK7-negative (PTK7^-^); thus PTK7^+^ CTCs represented less than 1% of all CTCs detected in the studied CRC patients (**Figure 2C**). PTK7 expression was detected more frequently on single CTCs than in CTC clusters (*data not shown*). The few patients (n = 7) with PTK7^+^ CTCs had primary tumors with an IHC score of 1 or 2 for PTK7 expression. None of the patients with primary tumors with an IHC score of 3 (n = 8, **Figure S2B**) had PTK7^+^ CTCs (**Figure S2C**). The presence of PTK7 on CTCs was therefore not limited to tumors with the highest PTK7 expression in the corresponding primary tumor.

**Figure 2:**
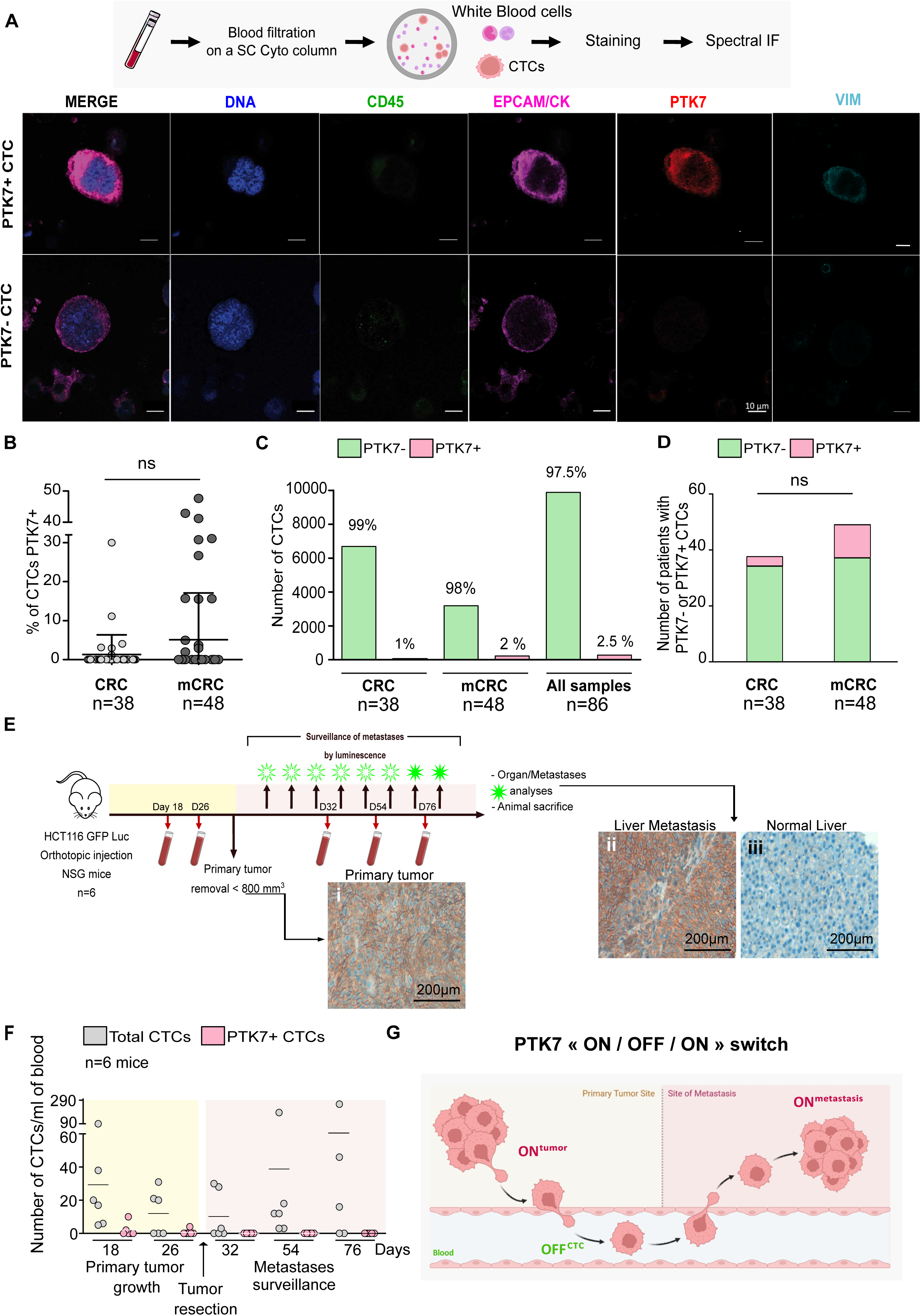
PTK7 is expressed frequently on primary and metastatic colon cancer cells but rarely on CTCs. **A.** Workflow recapitulating the steps to perform analyses of CTCs by immunofluorescence using the ScreenCell® (SC) Cyto columns (top) and representative images of immunofluorescence staining of CTCs isolated from the blood of CRC patients (bottom). CTCs are recognized by the expression of the epithelial markers EPCAM and/or pan-CK and/or VIMENTIN and by the absence of CD45. The expression of PTK7 was assessed on these cells. The upper images show a PTK7+ CTC and the lower images show a PTK7- CTC. **B.** Percentage of CTCs in the blood samples from patients with primary CRC (n=40) or mCRC (n=51) expressing PTK7. Each dot represents one patient. **C.** Numbers and percentages PTK7- (green bars) and PTK7+ (pink bars) CTCs in CRC patients with primary or/and metastatic disease **D**. Distribution and contingency analysis of PTK7- or PTK7+ CTCs in blood samples from patients with primary or metastatic CRC. **E**. Workflow of the mice model, with sample analysis timepoint used to test the ON/OFF/ON switch hypothesis and representative images of IHC for PTK7 performed on primary tumor (i), liver metastasis (ii) and normal liver (iii) tissues. **F**. Blood was collected on 18, 26, 32, 54- and 76-days post-engraftment of the HCT116 cell line and analyzed for the number of PTK7- and PTK7+ CTCs by flow cytometry. **G**. Graphical summary of the ON/OFF/ON switch that occurs in tumor cells during their journey to the metastatic niche.

To further investigate our initial hypothesis that PTK7 expression could serve as a marker for CTCs with a higher risk of metastasis, we specifically examined CTCs from patients whose CRC progressed to the metastatic stage within 2 years of enrollment in our study (n =7). These patients may have had more advanced disease or undetected metastases at diagnosis. However, none of these patients had PTK7^+^ CTCs (**Figure S2D**).

We also examined PTK7 expression in CTCs from patients with metastatic CRC (n=48 samples, mean: 12 CTCs/mL, range: 1 to 38 CTCs/mL) to assess whether PTK7 is a marker for the presence of metastases rather than a pre-metastatic marker. Among these 48 patients, 13 (27%) had PTK7^+^ CTCs (range: 2% to 47% of their total CTCs) (**Figure 2B**, mCRC), but only 2% (69 of 3,407) of the total CTCs identified in these patients expressed PTK7 on their surface (**Figure 2C**). A contingency analysis of the number of patients with PTK7^+^ CTCs did not show any difference between patients with CRC and those with mCRC (**Figure 2D**).

Altogether, although PTK7 is frequently expressed in primary (ON^tumor^) and metastatic (ON^metastasis^) CRC, it is not present on the cell surface (OFF^CTC^) of the vast majority of single CTCs and CTC clusters (> 97.5%), even in patients with advanced or overtly metastatic tumors.

### PTK7 is absent on the surface of CTCs collected from a xenograft mouse model of CRC that replicates the complete metastatic process

To study the dynamic associated with the PTK7 ON^tumor^/OFF^CTC^/ON^metastasis^ sequence evidenced in CRC patients, we used a mouse model that recapitulates the entire metastatic cascade of CRC and enables tracking of CRC cells, from the primary tumor to the metastatic sites, as well as CTCs. We generated primary tumors with high and homogeneous PTK7 expression by engrafting NSG mice (n = 6) with HCT116GFP-Luc cells, a human CRC line with high endogenous PTK7 levels engineered to express an ectopic GFP-Luc reporter. The grown primary tumors were removed and analyzed for PTK7 expression. Mice were then monitored for metastasis development by bioluminescence, while their blood was regularly collected and tested for PTK7 expression in CTCs (**Figure 2E**).

All mice developed metastases in at least three different organs (**Figure S2E**). All primary tumors (**Figure 2Ei**) and metastases (**Figure 2Eii**) expressed high levels of PTK7, whereas the surrounding normal tissue did not (**Figure 2Eiii**). CTCs in the mouse blood samples were defined as viable cells negative for H-2Kd (a mouse-specific marker) and positive for GFP and EPCAM (**Figure S2F**). All mice had CTCs (gray dots) before primary tumor resection, but only one mouse had some PTK7^+^ cells (pink dots), which accounted for 6.5% of the total CTCs examined, at 18 and 26 days after transplantation (**Figure 2F**). After resection of the primary tumors, all examined CTCs were PTK7^-^ (days 32, 54, and 76). Thus, by mimicking the metastatic process from the primary tumor to the development of metastases in a mouse model, we confirmed that PTK7 is not present on the surface of CTCs, even when both the primary and metastatic tumors were PTK7^+^. This confirms that PTK7 expression on epithelial tumor cells undergoes an ON^tumor^/OFF^CTC^/ON^metastasis^ switch during the metastatic process (**Figure 2G**).

### The PTK7 ON^tumor^/OFF^CTC^/ON^metastasis^ switch is a cell-autonomous process

To investigate whether the observed PTK7 ON^tumor^/OFF^CTC^/ON^metastasis^ switch could be influenced by the mechanical constraints experienced by CTCs, we characterized early plasma membrane remodeling induced by fluidic stress. Surfaceome proteomic profiling of HCT116 CRC cells was performed under three conditions: non-injected adherent cells, cells exposed to a 4-hour fluidic circulation system (circulating cells), and cells allowed to re-adhere after circulation (re-adherent cells). This fluidic system reproduces key physical aspects of bloodstream-associated mechanical stress through controlled flow conditions (4 ml/min and 2 dyn.cm^-2^). The short circulation time was selected to capture early adaptive responses to shear stress while limiting proteomic changes, including cell death, that could confound interpretation of surface signaling dynamics. At this early time point, PTK7 levels at the cell surface were not yet significantly affected, consistent with the need for prolonged fluidic exposure to induce its downregulation (see below).

Compared to short-term shear exposure, non-injected adherent cells showed increased post-translational protein phosphorylation, non-canonical NF-κB signaling, MAPK6/K4 signaling, and negative regulation of NOTCH4 signaling (**Table S7**). Conversely, cadherin, immunoglobulin superfamily cell adhesion molecules (IgSF CAMs) and Eph-ephrin signaling were increased in circulating cells compared to non-injected adherent cells, consistent with a perturbation in the cell adhesion state in response to short-term shear exposure (**Figure S3A,** *left panel* AND **S3B**). The transition between circulating and re-adherent states also produced marked remodeling of cell surface signaling (**Figure S3C**). Compared to re-adherent cells, circulating cells displayed significant upregulation of Rho GTPase signaling and Eph-ephrin signaling, indicating activation of cytoskeletal remodeling and stress-adaptive transcriptional programs associated with suspension and mechanical stress. Circulating cells exhibited coordinated downregulation of Hippo, NOTCH3, and syndecan-mediated adhesion signaling pathways relative to re-adherent cells (**Figure S3A,** right panel). Taken together, these changes define a reversible circulating-cell state characterized by enrichment of Rho GTPase-associated signaling, cytoskeletal activation and potential loss of stable epithelial and adhesion programs, which are restored upon re-adhesion.

To further investigate membrane and adhesion-associated changes at longer exposure times (24h and 48h) in response to impaired attachment and mechanical stress, we then cultured PTK7^+^ HCT116 and SW480 cells in 2D conditions that promote cell-dish adhesion (**Figure S3D**), or as 3D cultures under ultra-low adhesion (ULA) conditions that impair cell-dish adhesion and cell-cell contacts. In both cases, the cells were cultured either in Ca^2+^-containing normal medium, which favors cell adhesion, or in Ca^2+^-depleted medium, which impairs E-cadherin-mediated cell-cell and cell-matrix interactions. Additionally, cells were exposed to a flow (**Figure 3A**). We analyzed all cell samples for PTK7 expression by flow cytometry (FACS) after 24 and 48 hours of culture, using cells cultured under the adhesion condition in Ca^2+^-containing medium as the control. Consistent with data observed in xenografted HCT116 cells (**Figure 2Ei**), all cells expressed high levels of PTK7 in the control condition (**Figure 3B-C**, purple histogram). Except for cells grown under adhesion condition in Ca^2+^-depleted medium, which did not significantly differ from control cells, cells cultured under other conditions had lower proportion of PTK7^+^ cells. Notably, compared to control cells, a smaller fraction of cells cultured under ULA condition expressed PTK7, with significant differences observed after 24 hours in Ca^2+^-depleted medium and after 48 hours in Ca^2+^-containing medium (**Figure 3B-C**, green and yellow histograms). In **Figure 3D**, although the percentage of total viable cells (dashed line) and the number of PTK7^+^ cells (green) decreased with increasing culture duration in the Ca^2+^-depleted ULA condition, the percentage of PTK7^-^ cells (turquoise) increased until the fourth day. Measurement of PTK7 expression combined with a cell death marker (Livedead aqua™) revealed that PTK7^+^ cells grown under Ca^2+^-containing ULA condition either rapidly lose PTK7 expression and remain alive (blue arrow), or die, leading to a gradual loss of PTK7 from the cell surface (red arrow) (**Figure 3E**). However, the strongest downregulation of PTK7 was observed when HCT116 cells were exposed to a flow (**Figure 3A-C**, orange bars and graph). In this condition, almost half of the viable cells had lost cell surface PTK7 expression after 24 hours, and only 30% of viable cells still expressed cell surface PTK7 after 48 hours (**Figure 3B**). Altogether, these results suggest that the loss of PTK7 from the cell surface may provide a survival advantage to malignant cells in non-adherent conditions. Similar results were obtained with the SW480 cell line (**Figure S3E-F**). To ensure that flow does not generally affect all adhesion molecules, we examined the expression of CD44v6, a marker of CTCs, and found that its expression measured by FACS and immunofluorescence was not affected under any of the above conditions (**Figure 3F, 3G**). Similarly, we looked at the expression of EPCAM, an adhesion molecule known to be downregulated during the EMT process, but still expressed on (some) CTCs. We found that EPCAM was differentially affected under the tested conditions. Indeed, all cells grown under non-adherent conditions, including those exposed to a flow, remained positive for EPCAM, although at a lower level than under control condition (**Figure S3G-H**). Thus, the expression of two CTCs markers, *i.e.* CD44v6 and EPCAM, was not regulated by the same mechanism that controls PTK7 expression at the surface of tumor cells exposed to flow conditions.

**Figure 3:**
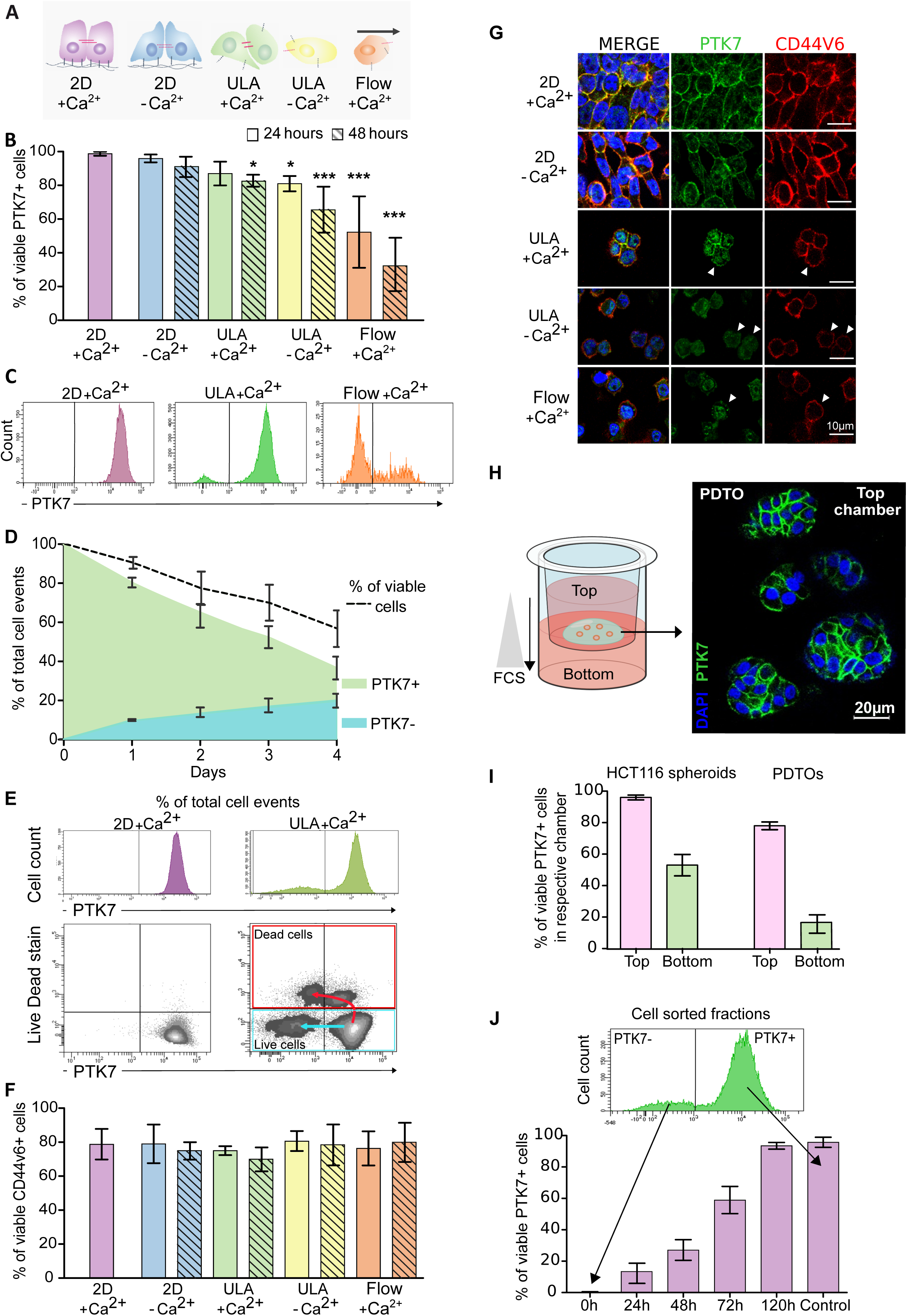
PTK7 ON/OFF/ON switch is triggered by the modification of cell-cell and cell-matrix contacts. **A.** Schematic representation of the different culture conditions, related to cell adhesion conditions, used in the study. HCT116 cells were cultured under classical 2D conditions in conventional cell culture dishes (2D) or in ultra-low adhesion dishes (ULA), with regular Ca^2+^ containing medium (+Ca^2+^) or Ca^2+^- depleted medium (-Ca^2+^) media. The cells were also exposed to a constant flow (flow +Ca^2+^). The color code of the different culture conditions is conserved throughout all figures. After 24 or 48 hours of culture, the cells were collected, stained and analyzed by FACS. **B.** Percentage of HCT116 cells expressing PTK7 grown in the indicated conditions of adhesion. An ANOVA with a Bonferroni post-tests was used to compare conditions. **C**. Representative flow cytometry histograms for the expression of PTK7 on HCT116 grown in the indicated condition. **D**. Percentage of viable HCT116 cells grown for 4 days under ULA condition with Ca^2^+ medium (dashed line). This percentage was calculated from the sum of PTK7+ (in green) or PTK7- (in blue) viable (live/dead™ negative) cells. **E**. Representative flow cytometry dot plots showing simultaneously the expression of PTK7 and the viability of HCT116 cell line grown in indicated conditions. The meaning of the blue and red arrows in the bottom right plot is explained in the main text of the result section. **F.** Bar graph showing the expression of CD44v6 on HCT116 cells exposed to the indicated adhesion conditions. Statistical analysis is similar to **B**. **G.** Representative images of immunofluorescence staining of PTK7 (green), CD44v6 (red) and DAPI (blue) on HCT116 cells grown for 48 hours in indicated conditions. **H.** Spontaneous release of tumor cells from tumor spheroids. *Left*: schematic representation of the Boyden chamber experiment. The insert (top part) containing medium without serum and HCT116 spheroids or PDTOs is placed in an ULA plate. Chemo-attractive medium (20% serum) was put in the bottom part of the ULA plate. *Right*: a representative image of immunofluorescence staining of PDTOs growing in the upper part of the Boyden chamber (DAPI in blue and PTK7 in green). Scale bar: 20µm. **I**. Bar graph showing the percentage of PTK7+ cells observed in the top and the bottom chamber after four days of culture. Mean value ± SD was calculated from n = 3 experiments with HCT116 spheroids and n = 3 experiments with PDTOs. **J.** Recovery of PTK7 expression after cell sorting of PTK7- cells. HCT116 cells were cultured in ULA +Ca^2+^ for 48 hours then sorted into PTK7- and PTK7+ populations by FACS (top). The sorted PTK7- populations was re-plated in adherent 2D +Ca^2+^ condition. PTK7 expression was then measured by FACS at the indicated time after plating and expressed as the number of PTK7+ cells in the whole population and compared to the control condition. For all experiments, results are expressed as the mean of at least three independent experiments ± SD and p-values were determined using a two-way ANOVA (*p <0.05, **p <0.005; ***p < 0.0005).

We then used a cell migration assay that mimics the spontaneous detachment of cancer cells from a tissue, and examined PTK7 expression in both adherent cells and cells that have migrated. To this end, we placed PTK7^+^ HCT116 spheroids embedded in Matrigel™ in the upper chamber of a Boyden chamber and induced migration to the lower ULA chamber using fetal bovine serum as a chemoattractant (**Figure 3H**, *left panel*). After 72 hours, we collected the contents of the chambers, and analyzed PTK7 expression on HCT116 spheroids in the upper chamber by immunofluorescence and on HCT116 single cells in the lower chamber by FACS. All cells in the upper chamber expressed PTK7 (**Figure S3I**); however, only 53% of the cells in the lower chamber still expressed PTK7 (**Figure 3I**). Similar results were obtained using patient-derived tumor organoids (PDTOs) (n = 3 CRC patients). Most cells in PDTOs placed in the upper chamber expressed PTK7 at the start of the experiment (**Figure 3H**, *right panel*) whereas only 20% of viable cells that migrated to the lower chamber still expressed PTK7 (**Figure 3I**, *bars on the right*).

Overall, our *in vivo* and *in vitro* data confirm the PTK7 ON^tumor^/OFF^CTC^ switch observed in CRC patients, and show that loss of cell-matrix and cell-cell adhesion leads to downregulation of PTK7 from the cell surface of CRC cells. The greater the loss of contact is, the higher the proportion of epithelial cells negative for PTK7 is observed.

Next, we investigated the extinction of PTK7 expression as a reversible phenomenon in the context of the ON^tumor^/OFF^CTC^/ON^metastasis^ switch. To this end, we harvested HCT116 cells exposed to a flow for 48 hours and re-plated them under fully adherent conditions. After 48 hours, all cells adhered to the plate and re-expressed PTK7 on their surface at a level comparable to that of the control condition (**Figure S3J**). Since the population obtained after fluidic exposure did not consist solely of PTK7^-^ cells, it remains possible that the growth of the few remaining PTK7^+^ cells exceeded that of PTK7^-^ cells. We therefore generated and sorted PTK7^-^ and PTK7^+^ cells, and plated them separately under fully adherent conditions for 5 days. All cells adhered to the plate. PTK7 expression on the cell surface was detected as early as 24 hours, increased progressively, and reached >95% of the population by the fifth day, similar to the control cells (**Figure 3J**).

Taken together, these *in vitro* experiments parallel the *in vivo* observations and confirm that changes in cell adhesion conditions trigger a versatile ON^tumor^/OFF^CTC^/ON^metastasis^ switch that controls, at least in part, the expression of PTK7, but not of CD44v6 or EPCAM, on the surface of epithelial tumor cells. The loss of PTK7 progresses over time when adhesion conditions are not maintained, and occurs rapidly in cells exposed to flow condition. Cells that adopt the PTK7^-^ phenotype appear to have a survival advantage over cells that remain PTK7^+^ under non-adherent conditions, at least for few hours. Moreover, the loss of PTK7 is reversible upon re-establishment of cell adhesion and cell-cell contacts.

### The PTK7 ON^tumor^/OFF^CTC^ switch is associated with increased cell survival and YAP1 activation in CTCs

To investigate the biological differences behind the PTK7^-^ cells and PTK7^+^ cells when adhesion is lost, we analyzed the signaling pathways of PTK7^-^ and PTK7^+^ cells. For this, we compared the transcriptome of individual viable PTK7^+^ (n=21) and that of PTK7^-^(n=14) HCT116 cells cultured under ULA condition (**Figure S4A**). The clustering of the 1,284 most variable genes showed that 208 genes were more highly expressed in PTK7^-^cells (Cluster 1), while 1,076 genes were more highly expressed in PTK7^+^ cells (Cluster 2) (**Figure 4A**, **Table S8**).

**Figure 4:**
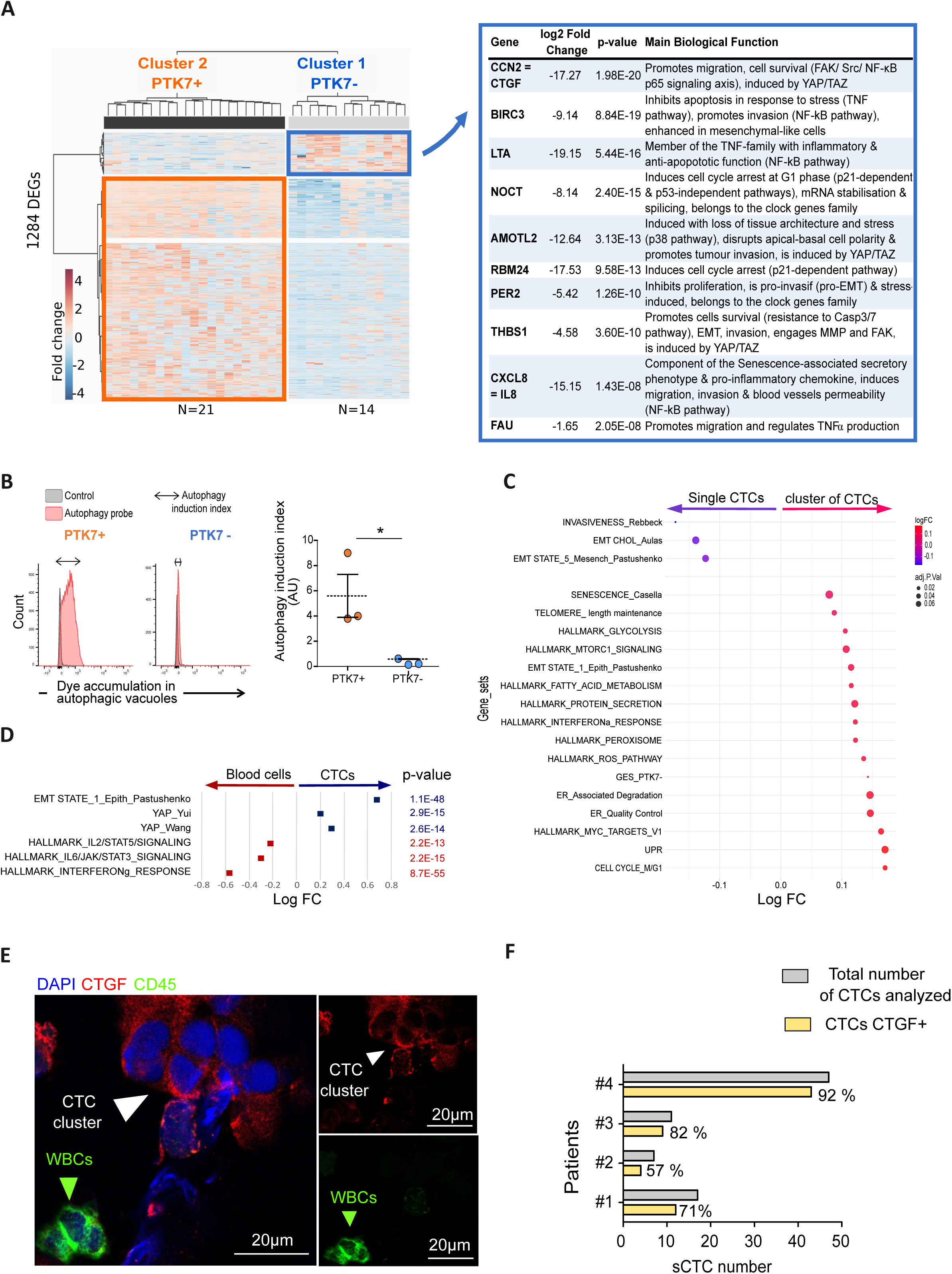
Single cell analysis shows that PTK7- cells escape anoikis-mediated cell death by activating the YAP1 signaling pathway and exhibit higher metastatic efficiency. **A**. RNAseq profiles in single HCT116 PTK7+ or PTK7- cells. Hierarchical clustering of PTK7+ (n = 21) and PTK7- (n = 14) single cells (left panel) and top 10 genes upregulated in PTK7- single cells compared to PTK7+ single cells (right panel). **B**. Autophagy induction in HCT116 PTK7+ and PTK7- cells. *Graph on the left:* Flow cytometry histograms showing autophagic activity in PTK7+ and PTK7- HCT116 cells after induction of autophagy with rapamycin (autophagy induction index is represented by the double arrow). *Graph on the right*: Autophagy induction index assessed in three independent experiments. **C.** Bubble plot representation of significant functional hallmark GSEA metagenes and relevant signatures enrichment in single CTCs from cancer patients (low survival ability in the blood flow) compared to cluster (higher survival ability in the blood flow compared to single CTCs) of CTCs from cancer patients. **D.** Representation of the top 3 functional hallmark GSEA metagenes and relevant signatures enrichment in all type of CTCs (single + cluster) from cancer patients compared to blood cells. **E**. Confocal analysis of CTGF (cytoplasmic and membranous, in red), CD45 (membranous, in green), DAPI (nuclear, in blue), in a CTC cluster (indicated with the white arrow head) of patients with CRC (CTC-colon Cohort). The white blood cell (WBC, green arrow head), is an internal negative control for CTGF staining. **F.** Bar plot showing the number of CTCs expressing CTGF (yellow bars) relative to all CTCs (grey bars) in n = 4 CRC patients. Data are expressed as mean +/- SD. P-values were determined using a one-tailed non-parametric t-test (*p <0.05).

The top 10 genes expressed in PTK7^-^ cells encode key factors involved in cell migration cell survival in response to stress and cell-cycle arrest, and that are regulated by TNFα, NF-κB and the YAP/TAZ signaling pathways. Ontology analysis using the KEGG and Reactome signaling pathways revealed activation of the “NF-κB pathway”, “Senescence-Associated Secretory Phenotype (SASP) and cellular senescence”, “Cellular responses to stress” and “Cytokine signaling” in PTK7- cells (**Table S8**). We also searched for enrichment of metagenes using the 50 GSEA hallmark gene sets (**Figure S4B,** upper radar plot). Among the metagenes tested, PTK7^-^ cells were enriched in metagenes that regulate inflammation (TNFα signaling *via* NF-κB and inflammatory response), response to stress (hypoxia, cell survival, UPR, senescence) and tumorigenic status (EMT, stemness score, cell cycle arrest in G1 phase, MYC targets and YAP signaling). To confirm activation of the YAP1 signaling pathway, we looked at CTGF expression, a YAP/TEAD-induced gene and the most highly expressed gene from the PTK7^-^ GES, in HCT116 cells recovered post-fluidic exposure by immunofluorescence. We observed an inverse correlation between CCN2/CTGF and PTK7 levels in HCT116 cells exposed to fluidic condition (**Figure S4C**), confirming the RNAseq data and supporting a change of phenotype in response to this experimental condition.

In parallel, we examined genes that were more highly expressed in PTK7^+^ cells (or downregulated in PTK7^-^ cells). The 10 most upregulated genes were involved in cell cycle regulation, suppression of EMT, and active metabolic and protein degradation pathways (*via* lysosomal activity) (**Table S9**). Consistent with this, gene ontology analysis revealed the activation of metabolic pathways, DNA damage, DNA repair, and lysosomal activities. Surprisingly, although numerous genes were upregulated in Cluster 2, fewer signaling pathways were identified than in Cluster 1. We analyzed metagene enrichment and found that PTK7^+^ cells exhibited a Type-I interferon (IFNγ and IFNα) response, a metabolic response (*e.g.* as bile acid, xenobiotic, and estrogen response), epithelial differentiation (intestinal differentiation, apical junctions) and autophagy-related pathways activation (*e.g.* glycolysis, lysosome, quiescence, macro-autophagy) (**Figure S4B,** *bottom*). To confirm this finding, we evaluated the activation of autophagy, an important survival mechanism under stress, in PTK7^+^ or PTK7^-^ HCT116 cells grown under ULA conditions. We monitored the accumulation of autophagic vacuoles after autophagy induction using a CYTO-ID® probe. We observed that PTK7^+^ cells, but not PTK7^-^ cells, exhibited autophagy flux (autophagy induction index of 5.6 +/- 2.94 in PTK7^+^ and 0.58 +/-0.94 in PTK7^-^ cells) (**Figure 4B**). This confirms that PTK7^-^ cells, unlike PTK7^+^ cells, do not trigger autophagy in response to a stress such as loss of cell adhesion.

To assess whether some of these findings were relevant in CTCs from patients, we tested metagenes related to some of these functions in our database. We found that single CTCs exhibited a more advanced mesenchymal state, while clusters of CTCs, which survive better in the bloodstream, were highly enriched with cells in the G1 phase and senescent state, as well as ER/peroxisome, MYC, mTOR, and interferon responses, and most importantly, with the PTK7^-^ GES (**Figure 4C**). Furthermore, the YAP1 signaling pathway was strongly activated in all CTC subsets (both individual CTCs and CTC clusters) compared to blood cells, as evidenced by two independent YAP1 signatures, notably the YAP1 signature from Yui *et al* (27). that relies on the upregulation of *CYR61* and *CTGF* genes (**Figure 4D**). We confirmed activation of the YAP1 pathway in CTCs by assessing the surface expression of CTGF in CTCs (**Figure 4E**). We found that 68 of 82 CTCs (88%) from CRC patients exhibit this expression pattern (**Figure 4F**), consistent with the activation of YAP1.

### The PTK7 ON^tumor^/OFF^CTC^ switch requires combined transcriptional and post-translational programs

We next investigated the possible mechanisms leading to PTK7 loss on CTCs and in tumor cells exposed to impaired adherent conditions. Loss of PTK7 expression from the cell surface can result from transcriptional regulation, endocytosis (+/- degradation), or shedding (28,29).

To investigate a possible transcriptional regulation, we first compared *PTK7* mRNA levels in HCT116 cells grown under adherent condition, ULA condition, or exposed to a flow. Similar to PTK7 protein levels, the *PTK7* mRNA levels decreased when cells were cultured for 48 hours under ULA condition with Ca^2+^-depleted medium or under flow condition in the presence of Ca^2+^ (**Figure 5A, Figure S5A**). Notably, mRNA levels remained “high” after 48h under flow condition, even though more than 70% of the viable cells were PTK7^-^ at the protein level. This statement is based on a comparison with WiDr, a CRC cell line used as a reference for low to intermediate PTK7 levels (**Figure S5B**). In parallel, we examined publicly available RNA sequencing data from CTCs and found, as previously shown (30), that most white blood cells (WBCs) have little or no detectable *PTK7* mRNA. Significantly more *PTK7* transcripts were observed in individual CTCs compared to leukocytes, although these levels were still below those observed in tumors. However, clusters of CTCs exhibited an intermediate amount of *PTK7* transcripts, which was not significantly different from the levels found in tumors and metastases (**Figure 5B**). These data suggest that the PTK7 ON^tumor^/OFF^CTC^ switch may depend in part on *PTK7* transcriptional regulation *in vitro* and in CTCs, but less clearly in CTC clusters from patients that were however PTK7^-^.

**Figure 5:**
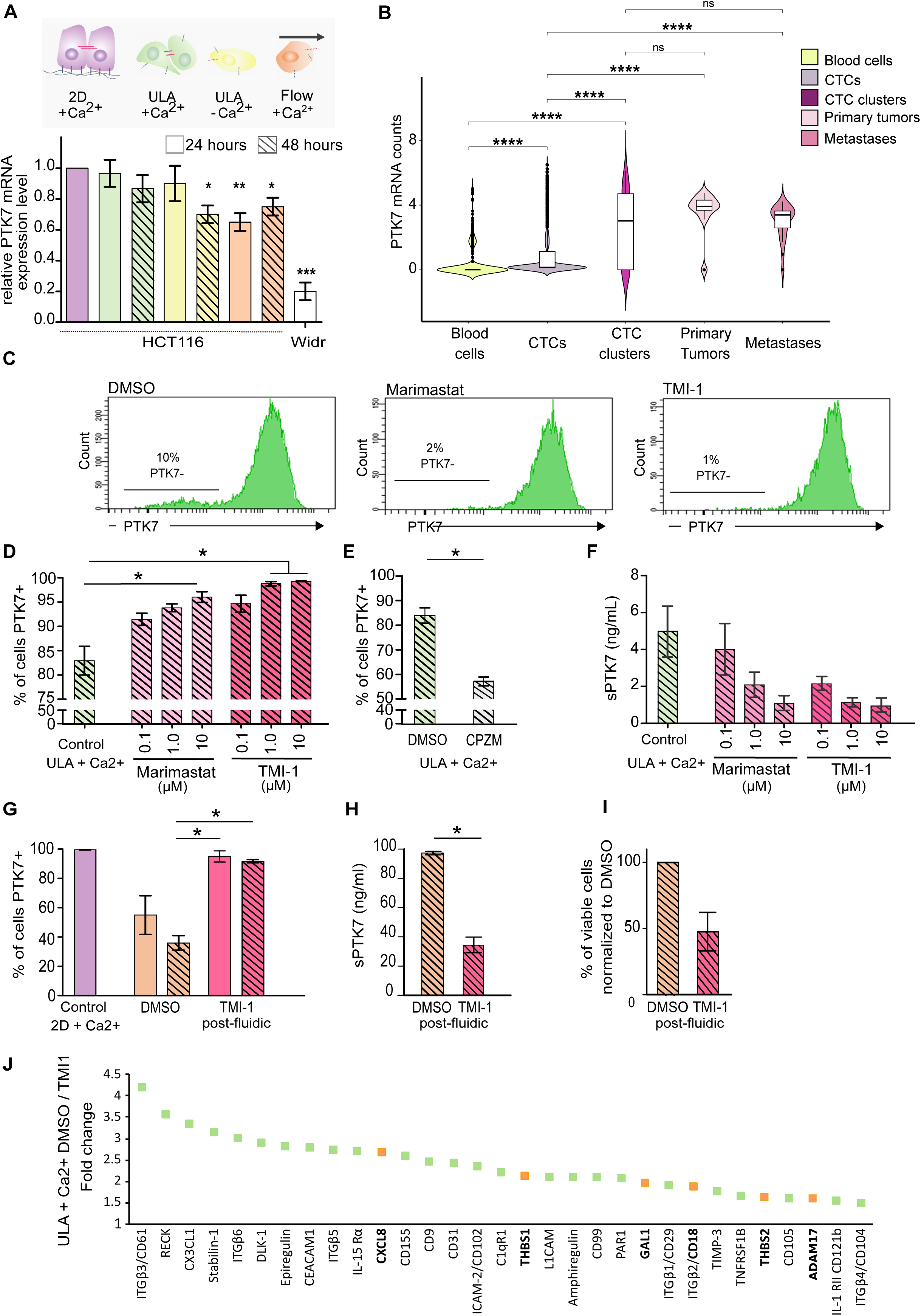
PTK7 ON/OFF switch involves the activation of MT1-MMP and ADAM17. **A.** Schematic representation of HCT116 cell culture conditions (Top) and relative *PTK7* mRNA levels, measured by RT-PCR, in HCT116 cells cultured in the indicated conditions for 24 and 48 hours (bottom). **B.** *PTK7* expression in CTCs (single CTCs and clusters of CTCs), compared to peripheral blood, primary or metastatic disease. The *PTK7* mRNA detection threshold was set to > 5 reads. **C**. Representative flow cytometry histograms of PTK7 expression after exposure to vehicle (left), marimastat (middle) and TMI-1 (right). HCT116 cells were cultured in ULA-Ca^2+^ medium and treated with marimastat or TMI-1 at a concentration of 10 µM for 48 hours. **D.** Percentage of PTK7+ HCT116 cells after exposure to increasing concentrations of marimastat and TMI-1 compared to cells cultured in ULA +Ca^2+^ medium and treated with DMSO (vehicle) for 48 hours. **E.** Percentage of PTK7+ HCT116 cells cultured in ULA +Ca^2+^ medium for 48 hours and treated for 4 hours with chlorpromazin or with DMSO (vehicle). **F.** Amount of sPTK7 (ng/ml) measured by ELISA in the supernatant of HCT116 cells treated under similar conditions as in (**D**). **G**. Percentage of PTK7+ HCT116 cells after submission to flow +Ca^2+^ and TMI-1 (10 µM) compared to cells exposed to flow +Ca^2+^ medium with DMSO (vehicle) and to the control condition (2D + Ca^2+^ medium) at 24 and 48 hours. **H.** Amount of sPTK7 (ng/ml) measured by ELISA in the culture medium of HCT116 cells treated under similar conditions as in (**G**). **I.** Viability of HCT116 cells exposed to flow for 48 hours, either with vehicle (DMSO) or with TMI-1 (10 µM) (similar conditions as in (**G**)). **J**. Results of the soluble receptor proteome profiler® array on the supernatant of HCT116 cells grown in ULA +/- TMI-1. Results are expressed as the fold change between these two conditions. Orange squares highlight molecules that were inversely affected by TMI-1 (inhibitor of MMP shedding activity) and chlorpromazine (facilitator of MMP shedding activity) exposition. Results are expressed as the mean of at least 3 independent experiments or as the mean of all patient samples ± SD. P-values were determined using a two-way ANOVA (*p <0.05, **p <0.005; ***p < 0.0005).

We thus explored other mechanisms that may be involved in reduction of PTK7 expression, such as endocytosis and cleavage, which are accessible to *in vitro* assays. PTK7 can undergo a caveolin-mediated endocytosis (31) or sequential cleavage by two matrix metalloproteinases (MMPs), MT1-MMP (MMP14) and ADAM metallopeptidase domain 17 (ADAM17 or TACE), leading to the shedding of its extracellular domain (soluble PTK7) (**Figure S5C**) (28,29). We assessed these mechanisms by adding inhibitors of caveolin-mediated endocytosis (Methyl-β-cyclodextrin) or MMP inhibitors (marimastat or TMI-1) to HCT116 cells cultured in ULA condition in the presence of Ca^2+^. No effect was observed with methyl-β-cyclodextrin (**Figure S5D**), whereas marimastat and TMI-1 treatments led to a dose-dependent increase in the percentage of PTK7^+^ cells compared to cells treated with the vehicle (DMSO) (**Figure 5C-D**). Similarly, treatment with chlorpromazine, an inhibitor of clathrin-mediated endocytosis that stabilizes MMPs at the cell surface (32), decreased the percentage of PTK7^+^ cells (**Figure 5E, S5E**).

To validate that MMPs indeed contribute to the PTK7^-^ phenotype, we measured the amount of shed soluble PTK7 (sPTK7) in the supernatant of cell culture using a custom ELISA. The concentration of sPTK7 was lower in the supernatant of cells treated with marimastat and TMI-1 compared to DMSO alone (**Figure 5F**), which inversely correlates with cell surface PTK7 expression (**Figure 5D**).

We then tested the involvement of MMPs activation in condition of flux. For this, we measured cell surface PTK7 expression in HCT116 cells under fluidic conditions, with or without TMI-1, and found that cells treated with TMI-1 had a higher percentage of PTK7 expression by FACS (**Figure 5G**) and lower amounts of sPTK7 were observed in the supernatant (**Figure 5H**). Most importantly, cells treated with TMI-1 in fluidic condition had lower survival than those treated with DMSO as the vehicle control (**Figure 5I**). Blocking the PTK7 ON^tumor^/OFF^CTC^ switch with a MMP inhibitor thus reduced the viability of cells exposed to a flow condition that mimic some of the cues, including impaired adhesion encountered in the bloodstream.

Shedding activity of PTK7 also occurs in the malignant context, notably we found higher levels of sPTK7 in the serum of patients with CRC (19 ng/mL) and mCRC (24ng/mL) than in healthy individuals (3.9 ng/mL) (p <0.0001 for both, **Figure S5F**). Interestingly, there was a trend toward a correlation between sPTK7 levels and the number of single CTCs counted (**Figure S5G**). Although we cannot be certain about the origin of the measured sPTK7, this finding suggests that PTK7 shedding from tumor cells is an important mechanism during the oncogenic process. Consistent with this, MMP14 and ADAM17 were also found in the serum of patients with CRC (**Figure S5H**). The use of specific inhibitors for MMP14 (NSC405020) and ADAM17 (GW280264X), revealed that after 48h of treatment, both GW280264X and NSC405020 showed a dose-response effect, with ADAM17 being primarily responsible for most PTK7 shedding in response to loss of adhesion in HCT116 cells (**Figure S5I**). To demonstrate the implication of active MMPs in PTK7 shedding, we briefly exposed the cells grown under ULA conditions to PMA, an activator of MMPs, +/- GW280264X. Results showed that PMA increased the number of PTK7^-^ cells in only 4 hours, but not when GW280264X was present (**Figure S5J**).

Since MMP14 and ADAM17 have numerous targets besides PTK7, we wondered whether other molecules expressed at the cell surface might be regulated by shedding activity. We therefore performed a soluble receptor proteome profiler array to investigate other potentially cleaved targets in the supernatant of HCT116 cells grown under ULA condition with or without TMI-1 (inhibitor of MMPs). Soluble factors and shed receptors were semi-quantified, using the control condition as the reference. In the presence of TMI-1, several factors were not shed (**Figure 5J**). A “reverse” experiment was performed using chlorpromazine (facilitator of MMP shedding activity) (**Figure S5K)**. Among the factors that were found decreased with the TMI-1 treatment compared to control, and increased with chlorpromazine compared to control, we found: CXCL8/IL-8, Nectin-2/CD112, THBS1, THBS2, GAL1, and ADAM17. Interestingly, two of these markers, CXCL8 and THBS1, were also found to be over-expressed by PTK7^-^ HCT116 cells grown in ULA condition and associated to YAP1 activation (**Figure 4A**).

In conclusion, these results suggest that the PTK7 ON^tumor^/OFF^CTC^ switch involves MMPs-mediated shedding that remodels the tumor cell surfaceome of malignant cells with CTC properties.

### The PTK7 ON^tumor^/OFF^CTC^ switch increases the metastatic potential of CRC cells

To investigate the effects of the PTK7^-^ phenotype on the metastatic potential of tumor cells, we prepared HCT116 cells grown under ULA condition and injected FACS-sorted PTK7^-^ and PTK7^+^ viable cells separately into the tail vein of mice. We then monitored the mice weekly for 4 weeks using bioluminescence imaging to detect metastases in each group (**Figure 6A**). A bioluminescence signal was observed in the group injected with PTK7^-^ cells as early as day 16 (p = 0.02, **Figure 6B**, statistics in blue), but not in the group injected with PTK7^+^ cells until day 23 (p = 0.049, **Figure 6B**, statistics in orange). The mean bioluminescence intensity was higher in the PTK7^-^ group than in the PTK7^+^ group at day 16 (p=0.02), day 20 (p=0.03), day 23 (p=0.049) and day 28 (p=0.046) (**Figure 6B**, statistics in black). After the mice were sacrificed on day 28, all organs were collected and analyzed by bioluminescence (**Figure 6C**). The total (**Figure S6A**) and average (**Figure 6D**) numbers of metastatic foci (detected by bioluminescence) were higher in mice injected with PTK7^-^ cells than in those injected with PTK7^+^ cells. The number of organs with detectable metastases was not statistically different, but tended to be higher in the group injected with PTK7^-^ cells (**Figures S6A, S6B**). These results suggest that PTK7^-^ cells may have better survival abilities in the blood leading to enhanced metastatic potential compared to PTK7^+^ cells.

**Figure 6:**
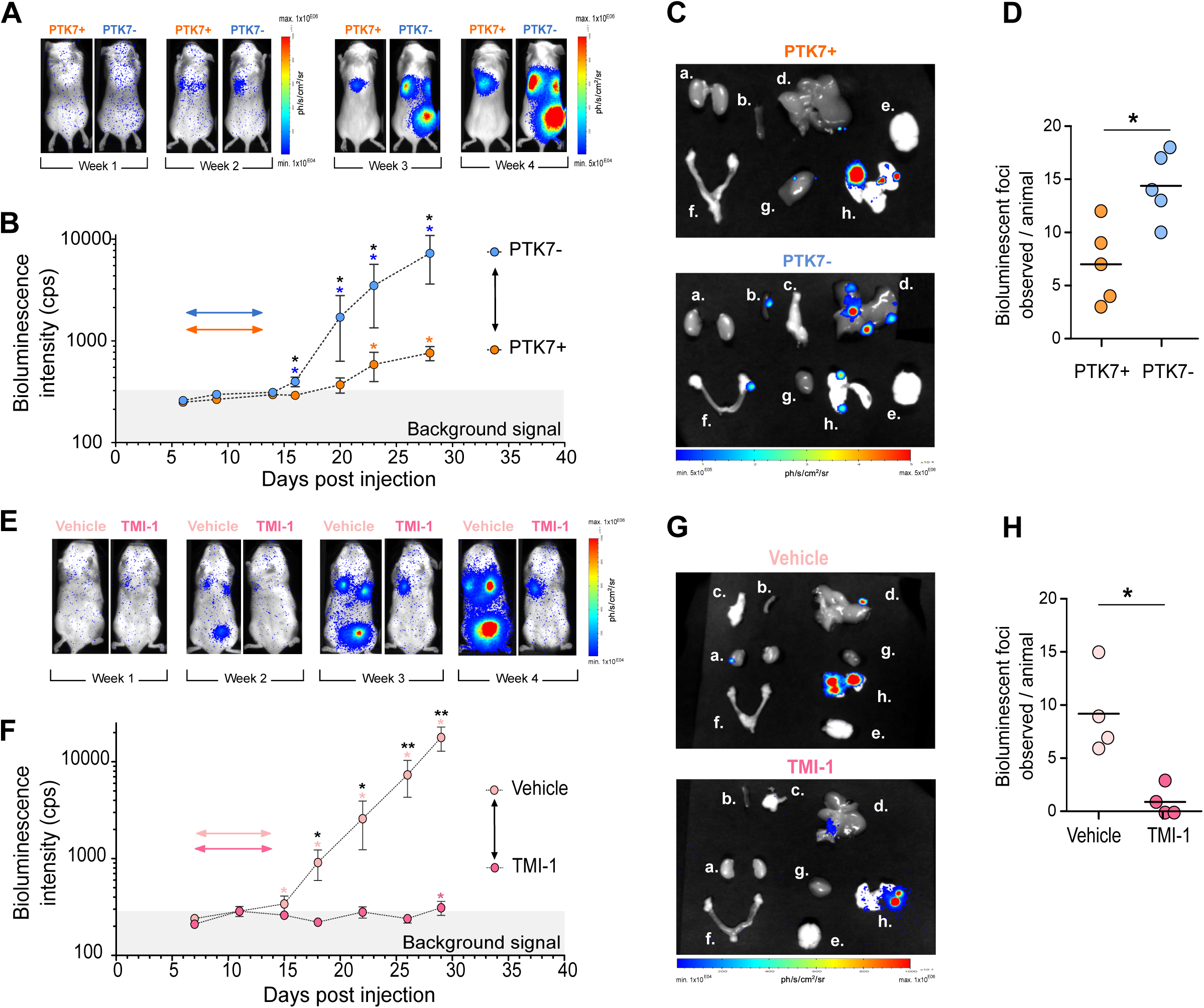
The shedding of PTK7 increases the metastatic potential *in vivo*. **A.** Representative bioluminescence images of NSG mice follow-up for metastatic occurrence after caudal injection of FACS-sorted PTK7+ (orange) or PTK7- (blue) HCT116 cells. **B.** Kinetics of bioluminescence intensity measured in mice injected with PTK7+ (orange) or PTK7- (blue) HCT116 cells (n = 5 mice per group). The statistics in blue and orange compare the signal intensity measured at the indicated time point with the value of reference, obtained 6 days after cell injection, in each group of mice (*ie.* injected with PTK7- cells or PTK7+ cells, respectively). The statistics in black compare the signal intensity observed at each time point between the two groups of mice (PTK7+ in orange dots and PTK7- in blue dots) groups. **C.** Example of bioluminescent metastatic foci located in different organs (a. Kidneys, b. spleen, c. pancreas, d. liver, e. brain, f. ovaries, g. heart, h. lungs) at autopsy of a mouse injected with the PTK7+ cells (top) and a mouse injected with the PTK7- cells (bottom). **D.** Average number of bioluminescent metastatic foci observed per mouse after injection of the PTK7+ (orange) versus the PTK7- (blue) HCT116-GFP-luc cells. **E.** Representative bioluminescence images of NSG mice follow-up for metastatic occurrence after caudal injection of HCT116 cells pre-treated with a TMI-1 or the vehicle. Prior to cells injection, mice were prepared with 2 injections of the *ad-hoc* treatment (TMI-1 or vehicle solution). **F.** Kinetics of bioluminescence intensity measured in mice injected with HCT116 cells treated with TMI-1 or the vehicle (n = 5 mice per group). The statistics in light pink and pink compare the signal intensity measured at the indicated time point with the value of reference, obtained 6 days after cell injection, in each group of mice (*ie.* injected with PTK7- cells or PTK7+ cells, respectively). The statistics in black compare the signal intensity observed at each time point between the two groups of mice (vehicle in light pink dots and TMI-1 in pink dots). **G.** Example of bioluminescent metastatic foci located in different organs (a. Kidneys, b. spleen, c. pancreas, d. liver, e. brain, f. ovaries, g. heart, h. lungs) at autopsy of a mouse injected with HCT116 cells pre-treated with TMI-1 (bottom) or vehicle (top). **H.** Average number of bioluminescent metastatic foci observed per mouse after injection of HCT116 cells pre-treated with TMI-1 or the vehicle.

In the absence of specific inhibitors of PTK7 cleavage, we next sought to evaluate how inhibition of MMPs-mediated might impact on metastasis occurrence. To this end, we pre-treated mice with TMI-1 (100 mg/kg) 24 hours and 1 hour before injecting HCT116-GFP-Luc PTK7^+^ cells (20,000 cells *per* injection), which had been pre-incubated with TMI-1 for 24 hours, into the tail vein. The control group was pre-treated with the vehicle and injected with 20,000 HCT116-GFP-Luc PTK7^+^ cells pre-incubated with the vehicle. The mice were monitored by bioluminescence for 4 weeks (**Figure 6E**). A bioluminescence signal was detected as early as day 15 in the vehicle-treated group (p = 0.049, **Figure 6F**, statistics in light pink) but not until day 29 in the TMI-1-treated group (p = 0.048, **Figure 6F**, statistics in pink). Additionally, the mean bioluminescence intensity was higher in the vehicle-treated group than in the TMI-1 treated group at day 18 (p = 0.030), day 22 (p = 0.029), day 26 (p=0.031) and day 29 (p=0.031) (**Figure 6F**, statistics in black). On day 30, all organs were collected from sacrificed mice and analyzed by bioluminescence (**Figure 6G**). Compared to the vehicle-treated cells, the TMI-1-treated cells formed significantly fewer metastases (n = 32*vs* 4, respectively) (**Figure 6H, S6A**). Accordingly, TMI-1 treatment also reduced the number of organs affected by metastases (7 in the control group and 2 in the TMI-1-treated mice) (**Figure S6A and S6C**).

Overall, we confirm that PTK7^-^ cells have a higher metastatic potential than PTK7^+^ cells. Importantly, inhibition of MMP-driven shedding of cell surface molecules including PTK7 by TMI-1 treatment reduces the metastatic potential.

## DISCUSSION

Our work shows that PTK7 expression, a receptor important in epithelial tissue organization, with functions in cell adhesion, migration and polarity, is an independent poor prognostic factor for overall survival in CRC patients. This prompted us to investigate PTK7 as a potential marker for CTCs at higher risk of metastasis. Against our expectation, PTK7 expression follows an ON^tumor^/OFF^CTC^/ON^metastasis^ sequence in tumor cells, during the metastatic process. We observed that the vast majority of CTCs from both human CRC patients and the mouse CRC model recapitulating the entire metastatic cascade do not express PTK7 on their cell surface (PTK7 OFF^CTC^, **Figure 7A**). Downregulation of other epithelial molecules, notably EPCAM and cytokeratins, is frequently observed in CTCs, in response to EMT (33–35), but not PTK7 (36). We thus assumed that, in this particular case, PTK7 loss in CTCs was driven by other mechanisms, and we investigated the PTK7 OFF^CTC^ phenotype specifically in the context of epithelial stress induced by the loss of adhesion, a situation encountered by CTCs in the blood flow, using several *in vitro* and *in vivo* models. Notably, we wondered if the modulation of this cell surface molecule in this context may play a role in cell survival and metastatic dissemination (37).

**Figure 7:**
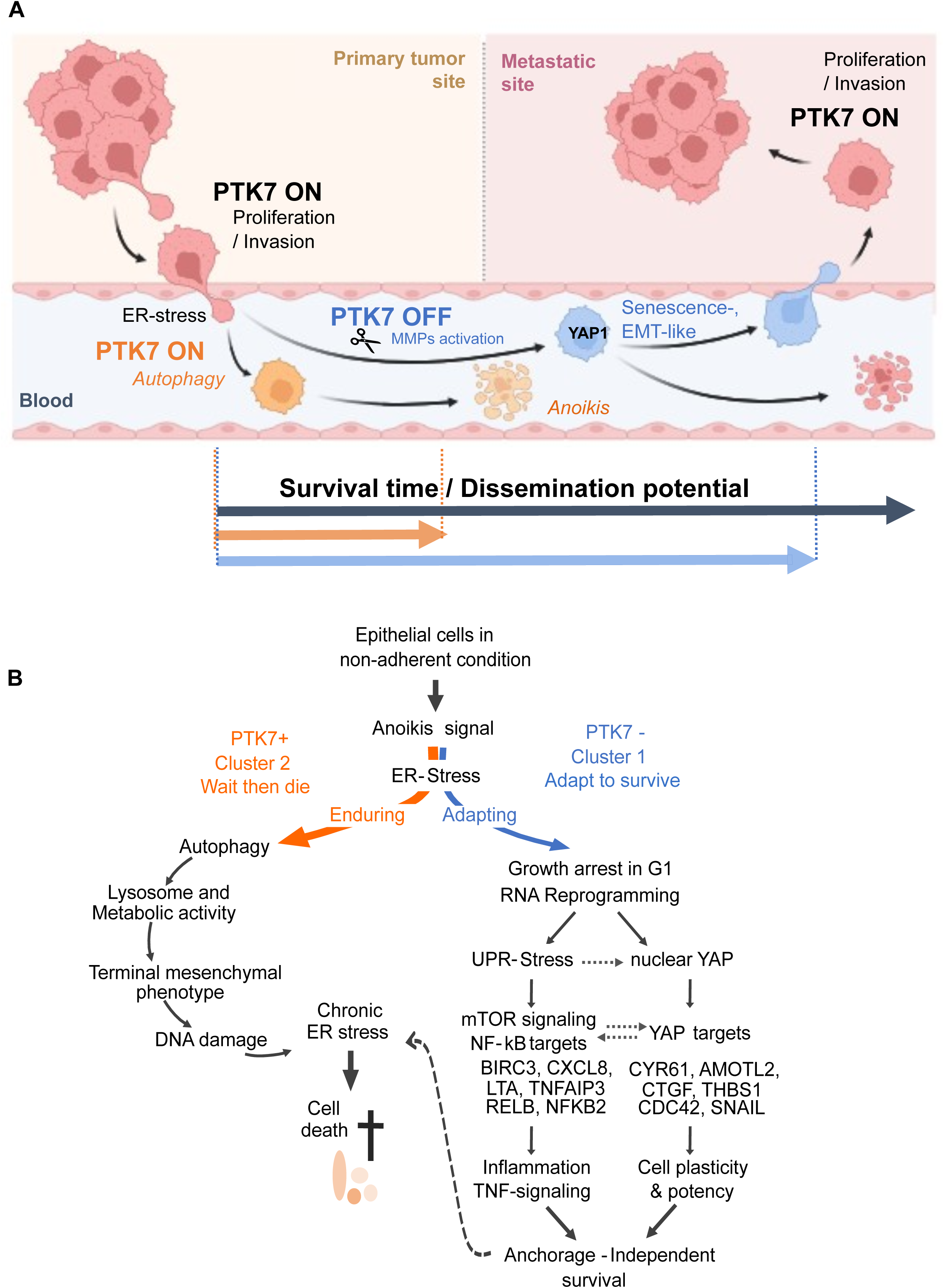
Summary of the pathways activated in CTCs based on PTK7 sequential expression. **A.** Graphical summary of the correlation between the phenotype of tumor cells (PTK7 ON/OFF/ON switch) and their survival and dissemination potential in the bloodstream. **B**. Graphical summary of gene ontologies associated with the PTK7+ or PTK7- cells.

### PTK7 OFF^CTC^ exhibit YAP1-mediated pro-survival signaling, EMT induction, and a senescence-like program

PTK7 is a catalytically inactive tyrosine kinase receptor that can propagate multiple signaling pathways depending on the presence and availability of co-receptors, ligands, and proteases at the plasma membrane (24,38). Full-length PTK7 activates the AKT and JUN pathways, promoting survival, cell proliferation, and migration (39). We did not observe enrichment in the AKT/JUN pathway in PTK7^OFF^ cells. However, PTK7 can also be internalized by endocytosis, and degraded (31), or cleaved and shed from the cell surface of cancer cells (40). In this case, the cleaved intracellular domain of PTK7, after nuclear translocation, can promote cell proliferation, migration and anchorage-independent colony formation through RAS/ERK and CREB/ATF1 signaling (39). In parallel, the N-terminal domain of PTK7 (sPTK7) activates non-canonical WNT/Ca2^+^ signaling through Frizzled receptors in epithelial cells. This paracrine mechanism leads to nuclear translocation of YAP1 and expression of YAP1/TEAD target genes (40). In our *in vitro* model and in CTCs from patients, we indeed observed significant activation of the YAP1 pathway, which can subsequently trigger the expression of key factors of cell plasticity and potency.

In our conditions, YAP1 activation was evidenced by the expression of key YAP1/TEAD target genes, notably *THBS1, CYR61, CDC42, SNAIL, AMOTL2,* and *CTGF/CCN2*. Interestingly, newly synthesized CTGF/CCN2 can have multiple autocrine functions. It can bind i) to the EGF receptor and β1 integrin, which then activate survival kinases such as AKT1, ERK1/2, SRC, and focal adhesion kinase (41,42); or ii) to TGFβ, forming a complex with high affinity for TGFβR, leading to EMT induction (43,44). We observed concomitant increased survival and EMT gene expression signature enrichment in our PTK7^-^ HCT116 cells and in CTCs.

YAP1 is also known to inhibit autophagy (7), and instead induces a senescence-like state (45). As a matter of fact, genes involved in the NF-kB pathway, which controls both cell-autonomous and non-cell-autonomous senescence programs, as well as IL8, a major SASP factor, were among the most highly activated genes in PTK7^-^ HCT116 cells grown under non-adherent conditions. The senescence-like metagene was also enriched in clusters of CTCs, which have better survival ability than single CTCs. Senescence deserves to be explored as a potential mechanism that may favor epithelial tumor cell survival under extreme stress conditions, notably in the bloodstream (46).

Altogether, this suggests that the sPTK7-mediated pathway may be favored in cells experiencing impaired adhesion and in CTCs, and that the YAP1/TEAD pathway is central to explaining the biology of PTK7^OFF^ cells.

Based on our combined *in vitro* and *in silico* transcriptomic data, we propose that tumor cells exposed to non-adherent conditions or CTCs in the bloodstream have one of two fates (**Figure 7B**). PTK7^ON^ cells activate the lysosome/autophagy signaling pathway to survive a brief period of stress (“Enduring” phenotype). However, these cells die rapidly if the stress persists. Alternatively, cells may acquire the PTK7^OFF^ phenotype, enter the G1 phase of the cell cycle, and initiate transcription of YAP1-TEAD target genes, such as CCN2/CTGF, to unlock the phenotypic plasticity of tumor cells (“adapting” PTK7^-^cluster).

Altogether, this suggests that the “adapting” PTK7^-^ cells may have a survival advantage over the “Enduring” PTK7^+^ phenotype.

### The PTK7 OFF^CTC^ involved ADAM17-dependent shedding of PTK7 in epithelial cells

The following question concerned how the dynamics and modulation of PTK7 expression occurs in response to impaired adhesion conditions.

We found that the PTK7 ON^tumor^/OFF^CTC^ switch occurs through combined transcriptional and post-translational processes following loss of cell adhesion. However, the transcriptomic downregulation was modest and insufficient to explain the PTK7 OFF phenotype observed in HCT116 cells, particularly under flow conditions reproduced *in vitro*, or in clusters of CTCs isolated from patients, where *PTK7* mRNA was still significantly present, and the protein absent. In patients, *PTK7* mRNA downregulation was more pronounced in single CTCs. This difference between single CTCs and CTC clusters could be related to residual cell-cell contacts in CTC clusters that may trigger basal *PTK7* expression, to the fact that CTC clusters extravasate more rapidly, and spend less time in the bloodstream than single CTCs, allowing less time for observable *PTK7* transcriptomic downregulation, or to the possibility that CTC clusters survive better in the blood flow than single CTCs. If *PTK7* downregulation is driven by loss of adhesion-induced stress, it will be thus less pronounced in single CTCs (47).

In our *in vitro* settings, the shedding of PTK7 was mainly driven by the metalloprotease ADAM17 (48,49). MMPs promote the cleavage of ECM components and receptors to regulate tumor cell functions during the metastatic cascade (50), such as migration (51,52), cell adhesion and signaling (53), and extra/intravasation (54). Interestingly, in CRC patients, a high level of serum MMPs is a marker for aggressive tumors with metastasis-promoting properties (28,55,56). Although the role of MMPs in tumor cells under adhesion conditions is well described (57), very little is known about what occurs when cells intravasate into the bloodstream or are in suspension. Recently, ADAM17 was shown to enhance the vascular permeability of tumor cells. ADAM17 is therefore active when tumor cells enter the bloodstream (58,59).

However, ADAM17 and MT1-MMPs have many other known targets, and their activity and preferred targets likely depend on substrate availability, which varies among tissues, during the extravasation phase, and in blood (60).

### Remodeling of the CRC cell surfaceome, including PTK7 shedding, makes cells more potent in promoting successful metastasis

We showed that under non-adherent conditions, MMPs are indeed active on targets other than PTK7, leading to broader remodeling of cell surface proteins (*e.g.,* Nectin-2), which may significantly impact downstream signaling. Interestingly, the shedding of various adhesion molecules may further contribute to the overall cell deformability and “long-term” resistance of CTCs to hemodynamic shear stress. Recent literature suggests that the ability of CTCs to deform and adapt to constricted vessels likely plays an important role in their survival, the circulation routes they take and the vascular sites where they arrest (61,62). Notably, our mouse model injected with PTK7- cells displayed more metastases in general, and more organs with metastases, suggesting that cells with a PTK7 OFF phenotype obtained after adhesion impairment are primed to resist bloodstream conditions and subsequent stresses encountered. This trait makes them more potent in promoting successful metastasis. In line with this, the therapeutic inhibition of PTK7 shedding, and probably of shedding activity in general, with a metalloprotease inhibitor impedes cancer progression and dramatically reduces the development of metastasis in mice. Despite the efficacy of MT1-MMP or YAP1/TEAD signaling pathway inhibitors in preclinical assays, numerous and serious side effects have limited the use of such treatments in patients to date (63,64). However, the PTK7 ON^tumor^/OFF^CTC^/ON^metastasis^ switch may offer new and interesting pharmacological opportunities, such as using CTGF inhibitors or senolytic agents, notably in combination with other treatments to target tumor cells at all stages of progression.

The discovery of the PTK7 ON^tumor^/OFF^CTC^/ON^metastasis^ sequence in CRC provides important insight into the mechanisms that govern CTCs survival and the development of metastasis occurrence in cancer patients (65,66). Nevertheless, this mechanism does not act in isolation, as multiple complementary processes also support CTC persistence in the bloodstream, as shown in breast and other cancer models (67–72).

Taken together, these findings strengthen our overall understanding of the metastatic cascade in patients and reveal a previously unappreciated, dynamic, and cell-autonomous regulatory mechanism that cancer cells engage upon entry into the circulation.

## METHODS

### Gene expression analysis of colon clinical samples

We gathered clinicopathological and gene expression data of tissues from normal colon (NC), primary colorectal adenocarcinoma samples (CRC) and metastases from CRC samples (mCRC) from 11 public data sets comprising at least one probe set representing *PTK7*. Sets and raw data were collected from the National Center for Biotechnology Information (NCBI)/Genbank GEO, ArrayExpress and TCGA databases (**Table S1**). Samples were profiled using whole-genome DNA microarrays (Affymetrix) or RNA sequencing (Illumina). The analyzed data set contained a total of 2,402 samples, including 95 NC samples, 2,239 CRC samples, and 68 mCRC samples.

Data analysis required a step of pre-analytic processing. We first normalized each data set separately, by using Robust Multichip Average (RMA) with the non-parametric quantile algorithm for the raw Affymetrix (73). Normalization was done in R using Bioconductor and associated packages. We then mapped hybridization probes across the different microarrays represented, as previously reported (74). When multiple probes mapped to the same GeneID, we retained the one with the highest variance in each data set. We log2-transformed the available TCGA RNAseq data that were already normalized. Next, we extracted *PTK7* mRNA expression and corrected the batch effects through the 11 studies using z-score normalization. Briefly, for each expression value in each study separately, *PTK7* values were transformed by subtracting the mean of the gene in that data set divided by its standard deviation, mean and standard deviation being measured on primary samples. *PTK7* expression was measured as a discrete value after comparison with median expression in the 95 NC samples; upregulation, thereafter designated “PTK7^High^” was defined by a CRC/NC ratio ≥1.5 and no upregulation (“PTK7^Low^”) by a CRC/NC ratio <1.5. The Consensus Molecular Subtype (CMS) classification was based on the tool CMScaller (75,76). Finally, to explore the biological pathways associated with our PTK7-based classification, we applied a supervised analysis using learning and validation sets. The learning set included the 459 samples of the TCGA data set, which included 92 PTK7^Low^ and 367 PTK7^High^ samples. We used a moderated t-test with empirical Bayes statistic included in the limma R packages. False discovery rate (FDR) was applied to correct the multiple testing hypothesis and significant genes were defined by the following thresholds: p <5%, q <25% and fold change (FC) superior to |2x|(Hochberg and Benjamini, 1990). A metagene score was derived from these 316 DEGs and was defined as the difference between mean expression of genes upregulated and mean expression of genes downregulated in the PTK7^High^ samples and using a cut-off equal to 0. This score was then applied to both learning and validation sets to test the robustness of the GES (t-test). Ontology analysis of the gene list was based on GO biological processes of the Database for Annotation, Visualization and Integrated Discovery (DAVID; http://david.abcc.ncifcrf.gov/).

Enrichment for several EMT Metagene-based signatures were tested: these metagenes were built from EMT Hallmark gene set (https://www.gsea-msigdb.org/gsea/msigdb/cards/HALLMARK_EPITHELIAL_MESENCHYMAL_TRANSITI ON.html), the MET and EMT/CHL signatures (77), the EMT tumor budding region signature (78), the l Core epithelial-to-mesenchymal transition interactome gene-expression signature (35), the intermediate states of EMT (79).

Regarding the statistical analyses, correlations between the *PTK7* expression-based classes (low *versus* high) and the clinicopathological factors were calculated with the Fisher’s exact test for the binary variables and the Student’s t-test for the continuous variables. Our primary endpoint, overall survival (OS), was calculated from the date of diagnosis until the date of death from CRC. The follow-up was measured from the date of diagnosis to the date of last news for event-free patients. Survival was calculated using the Kaplan-Meier method and curves were compared with the log-rank test. Univariate and multivariate analyses were done using Cox regression analysis (Wald test). The variables tested in univariate analysis included the PTK7-based classification (low *versus* high), patients’ age and sex, tumor location, pathological stage (based on the pTN staging) and grade, MMR status, and the CMS classification. Multivariate analysis incorporated all variables with a p-value inferior to 5% in univariate analysis. All statistical tests were two-sided at the 5% level of significance. Statistical analysis was done using the survival package (version 2.30) in the R software (version 2.15.2). The paper was written in accordance with reporting recommendations for tumor marker prognostic studies (REMARK) criteria (80).

### In vitro functional assay of EMT

HCT116 and SW620 cells were cultured under 2D conditions for 96 hours in DMEM with Ca2^+^, 10% FCS, 1% Hepes, and 1% P/S, with or without 1% StemXVivo (Ref: # CCM017, Bio-Techne®). Medium was renewed at 48 hours. Cells were then harvested, washed, and resuspended at 1 × 10^6 cells/ml in 1× PBS for subsequent flow cytometry analysis.

### Patients’ samples collection

Peripheral blood samples from subjects with primary (n=26) or metastatic (n=40) colorectal cancer were prospectively collected within the “CTC-colon” clinical trial. This unicentric trial was conducted at the Paoli-Calmettes Institute (Marseille, France), and registered as https://ClinicalTrials.gov identifier NCT03256084. It was proposed to patients with primary or metastatic colon cancers. Its primary objective was to provide a better understanding of which circulating tumor cells (CTCs) have the higher risk for metastasis, based on disease’s stage. The study was approved by the French National Agency for Medicines and Health Products Safety, a national ethics committee (CPP Sud-Méditerranée), and our Institutional Review Board. It was conducted in accordance to the Good Clinical Practice guidelines of the International Conference on Harmonization. The nature and possible consequences of the study were explained to the patients. All patients gave their informed consent for inclusion, biopsy and blood sampling, and molecular analysis.

Blood samples were collected in a 5mL Vacutainer® tube containing EDTA K2 as an anticoagulant. Before that, the first milliliters of blood were discarded to avoid endothelial cells contamination during the puncture. All samples were shipped to the laboratory, treated and analyzed within 4 hours.

Tissue from colorectal cancer patients from the IPC and the AP-HM hospitals, were collected after surgical resection in the framework of two clinical trials: primary tumors were collected in the frame of the CTC-colon trial, mentioned above, and liver metastases were collected in the frame of the B-Org cohort. The B-Org cohort was approved by the French National Agency for Medicines and Health Products Safety, a national ethics committee (CPP Sud-Méditerranée), the AP-HM Institutional Review Board (2019-A00710-57), and registered as https://ClinicalTrials.gov identifier NCT05384184. Again, informed consent was obtained after the nature and possible consequences of the study were explained. All samples were formalin fixed and paraffin embedded, and evaluated in the Biopathology department.

### Immunohistochemistry (IHC) analysis

Primary and metastatic tumors were fixed in formalin and embedded in paraffin for histopathological analysis and PTK7 staining. Section of 5□μm were submitted to a deparaffinization step in histolemon (Carlo Erba Reagenti, Rodano, Italy) (2 x 5 min). Sections were rehydrated in graded alcohol (100 % ethanol (5 min), 96% ethanol (3 min), 70% ethanol (3 min)) then, dH_2_O (5 min). For antigen enhancement, sections were incubated in target retrieval solution. Staining reactions were done at RT with an automatic stainer and were conducted as follows: after washes in phosphate buffer, followed by quenching of endogenous peroxidase activity with 3% H_2_O_2_, slides were first blocked with serum during 30 min and then incubated with the PTK7 antibody described in **Table S10**. After washes, slides were incubated with biotinylated antibody against goat immunoglobulin for 25 min and then by streptavidin-conjugated peroxidase. Diaminobenzidine was used as the chromogen. Hematoxylin was used for counterstaining, and coverslips were mounted using Permount™ media (Fisher Scientific). They were examined under a light microscope.

PTK7 staining was scored from 0 to 3 based on the staining intensity which is the most represented on the overall tumor cells. A score 0 corresponds to absence of staining; a score 1 corresponds to a light, diffuse and or apical staining; a score of 2 corresponds to a moderate staining (most often at the membrane level), and score 3 corresponds to a strong surface staining.

### Immunofluorescence on cell lines

Cells were seeded on coverslips 24h before the experiment. Coverslips were washed quickly with PBS before being fixed for 15 min with PBS 4% Paraformaldehyde (Thermo Scientific, Waltham, MA, USA). Cells were then permeabilized with PBS-0.5% TX100 (Euromedex, Souffelweyersheim, France) for 15min at room temperature. Coverslips were blocked with PBS-5% Normal Horse Serum (Sigma, Saint-Louis, MO, USA) for at least 30 min. Primary antibodies (**Table S10**) were diluted in blocking buffer and incubated 1 h at room temperature. Coverslips were washed three times for 5 min between primary and secondary antibody incubations. Subsequently, secondary antibodies (**Table S10**) were added along with DAPI for 1 h at RT. Cells were washed extensively and mounted with ProLong Antifade reagent (Invitrogen, Carlsbad, CA, USA). Pictures were taken with confocal microscope LEICA LSM880, 40X objective (LEICA, Wetzlar, Germany) using ZEN software.

### Cells lines, basic medium and reagents

Cell lines were purchased from the ATCC® collection and were regularly checked for mycoplasma contamination. SW480 cells (ATCC®CCL228™) were cultivated in DMEM (GIBCO) supplemented with 10% FBS (GIBCO), 1 mM HEPES (GIBCO). HCT116 cells (Horizon™) were cultured in RPMI (GIBCO) supplemented with 10 % FBS (GIBCO), 1mM HEPES (GIBCO), 1% Penicillin/Streptomycin (GIBCO).

### Staining of CTC by immunofluorescence

Three milliliters of blood were used with a ScreenCell® CYTO column (ScreenCell®, Sarcelles, France) to enriched CTC on the filter according to the manufacturer instructions. Immunofluorescent staining was then done directly on the filter. Cells were fixed using 4% paraformaldehyde for 5 min, permeabilized using TBS, 0.2% Triton X-100 for 5 min. After a rinse in water, filters were incubated in blocking buffer (TBS, 3% bovine serum albumin, 1% donkey serum, 1% goat serum) at room temperature (RT) for 30 min. Primary antibodies (listed in **Table S10**) were added in blocking buffer and incubated overnight (ON) at 4°C. Following 3 washes with TBS-0.05% Tween20, secondary antibodies were added (listed in **Table S10**) for 1 h in the dark at RT. After 3 washes with TBS-0.05% Tween20, the coupled antibody CD45-A488 (Biolegend, 304017) was incubated for 1 h in the dark at RT. Finally, after three washes with TBS-0.05% Tween20 and H_2_O, the filter was counterstained with Sytox Blue Nucleic Acid Stain for 5 min at RT in the dark to stain nuclei (Life Technologies, S11348). The filter was mounted with Kaiser Solution (Sigma Aldrich, 1092420100) and dried at RT a few hours. Immunofluorescence were analyzed with a c-Plan-Apochromat 40x/1.3 Oil objective on a LSM880 confocal with spectral detection from Zeiss equipped with a 405-laser diode, an Argon laser and 561 and 633 lasers. Imaged acquisition and spectra unmixing were done using the Zen Black software as described in Lopresti *et al* (81).

### Ethics statement for animal experiments

Studies on animals were conducted in accordance with the current ethical standards of the European Community (Directive 2010/63/EU), the Ethics Committee for Animal Experimentation (CEEA#14) and the French Ministry of Higher Education and Research, which approved and authorized the entire procedure described in this paper (project number APAFIS #35294).

### Xenograft mouse models

NOD-SCID IL-2Rγ null (NSG) mice were purchased from The Jackson Laboratory (NOD.Cg-*Prkdc^scid^ Il2rg^tm1Wjl^*/SzJ, Stock N° 005557). HCT116 cells were infected with lentivirus vector expressing GFP (Green fluorescent protein) and LUC2 (HCT116-GFP-Luc). 1.10^6^ cells were mixed in Matrigel™ (Corning) and subcutaneously injected into 8-week-old NSG mice. Tumor development was followed over a period of 4 weeks, and tumors were resected at a size of 750-1000 mm^3^. The development of metastases was monitored by bioluminescence analysis, using PhotonIMAGER^TM^ (BIOSPACELab). When metastatic signals appeared, the animals were euthanized and autopsied. The organs were analyzed for the presence of HCT116-GFP-Luc cells by bioluminescence. The images were analyzed with M3 Vision^TM^ (BIOSPACELab).

### Mouse blood sample collection

For studies of CTCs in mice, blood was obtained by retro-orbital sampling and up to 200 µL of blood was collected every 2 weeks in an EDTA-coated tube. Lysis of the red blood cells was performed with ACK lysis buffer (Gibco).

### Adhesion-impaired cell culture conditions

Cells in 2D culture were detached using TrypLe express solution (GIBCO), counted and dispensed in regular or in Ultra Low Attachment flask (Corning). Cells were suspended in the *ad hoc* culture medium, *i.e.* DMEM 10% FBS medium, either with or without Ca^2+^/Mg^2+^ for 24 or 48 hours

### *In vitro* fluidic assay

Cells in 2D culture were detached using TrypLe™ express solution (LifeTechnologies), or EDTA (LifeTechnologies), counted and suspended in culture medium containing 100mM Hepes (LifeTechnologies). Then cells were injected in a fluidic unit composed of: a perfusion set with 10mL reservoirs, a 0.8mm ID silicone tubing, an uncoated hydrophobic µ-slide I Luer channel of 0.8mm and an in-line Luer injection port (all from Ibidi Gmbh). Fluid circulation was provided by computer-controlled air pressure pump (Ibidi Gmbh), driven with the Ibidi PumpControl software v1.5.0. Cells were exposed to a low rate of 4mL/min for 24 or 48h.

### Patient-derived tumoroids

Patient-derived tumor organoids (PDTOs) were derived from patients with primary colorectal cancers as described previously (82).

### Transwell assay

A transwell insert (Corning®, 8.0 µm pore, # 3422) was placed in a 24 wells plate. BME Type 2 (Cultrex RGF Basement Membrane Extract, Type 2, # 3533-010-02) was layered on the membrane. PTDOs (≈10 000 PDTOs) were seeded on the upper transwell insert in PDTOs medium containing 5% BME. PDTOs medium supplemented with 20 % FBS was added to the lower chamber. An incubation at 37°C at 5% CO2 was performed for 2 days. PDTOs (upper compartment) and migrating cells (lower fraction) were collected and washed with 10 ml cold PBS1x. Upper fraction was used to perform immunofluorescence staining. Lower fraction was used for flow cytometry analysis.

### Flow cytometry analysis

After collection, cells were incubated 5 min with human BD Fc Block™ (BD Pharmigen™, 564220) at RT, to block non-specific antibody staining. Cells were then incubated with coupled primary antibody, described in **Table S10,** in FACS buffer and incubated for 30’ on ice. Samples were analyzed on a BD^TM^ LSRII flow cytometer (BD Biosciences). Data were then analyzed using the BD FACSDiva^TM^ and FlowJo^TM^ softwares.

### Single cell isolation

HCT116 cells cultured under ULA condition for 48 hours. Cells were harvested and stained with a cell death marker (Live dead orange) and an anti-PTK7-FITC (Mylteniy) according to manufacturer’s instructions. Viable single cells with the PTK7- or PTK7+ phenotypes were piked with a Transfer Man-K2 micromanipulator (Eppendorf™), transferred to a tube with lysis buffer plus polyA primers (one cell per tube) and frozen. The samples were then amplified using the Smart-seq™ library for single cells from Takara.

### Surfaceome analysis under circulating conditions

HCT116 cells were subjected to a fluidic circulation system for 4 h. Three conditions were analyzed: (i) non-injected adherent cells, (ii) circulating cells immediately collected after circulation, and (iii) circulating cells allowed to re-adhere for 2 h after circulation. Dead cells were removed using the STEMCELL Technologies EasySep Dead Cell Removal (Annexin V) Kit.

Cell surface biotinylation was performed as previously described by Guang-Ni Huang (83). Cells were washed twice with ice-cold PBS/2.5 mM CaCl_2_/1 mMMgCl_2_, pH 7.4 and incubated with 0.5 mg/ml Sulfo-NHS-SS-biotin on ice for 30 min. Unreacted biotin was quenched using 50 mM glycine in PBS/CaCl_2_/MgCl_2_. Cells were lysed in IP buffer (PBS containing 5 mM EDTA, 5mM EGTA, 10 mM sodium pyrophosphate, 50 mM NaF, 1 mM NaVO3, 1% Triton X-100, and protease inhibitors), and lysates were centrifuged at 14,000 x *g* for 10 min at 4 °C. Biotinylated proteins were isolated using immobilized NeutrAvidin beads for 2–3 h at 4 °C under rotation. Beads were washed three times with IP buffer, and proteins were eluted in 2× loading buffer at 65 °C for 10 min prior to mass spectrometry analysis.

### Mass spectrometry analysis

Peptides pulldown extracts were loaded on NuPAGE 4-12% Bis-Tris acrylamide gels (Life Technologies) to stack proteins in a single band that was stained with Imperial Blue (Thermo Fisher Scientific) and cut from the gel. Gels pieces were submitted to an in-gel trypsin digestion after cysteines reduction and alkylation (Shevchenko et al., 1996). Peptides were extracted from the gel and dried under vacuum. Samples were reconstituted with 0.1% trifluoroacetic acid in 4% acetonitrile and analyzed by liquid chromatography (LC)-tandem mass spectrometry (MS/MS) using an Orbitrap Fusion Lumos Tribrid Mass Spectrometer (Thermo Electron, San Jose, CA) both online with a nanoRSLC Ultimate 3000 chromatography system (Dionex, Sunnyvale, CA). Peptides were separated on a Thermo Scientific Acclaim PepMap RSLC C18 column (2µm, 100A, 75 µm x 50 cm). For peptide ionization in the EASY-Spray nanosource in front of the Orbitrap Fusion Lumos Tribrid Mass Spectrometer, spray voltage was set at 2.2 kV and the capillary temperature at 275 °C. The Orbitrap Lumos was used in data dependent mode to switch consistently between MS and MS/MS. Time between Masters Scans was set to 3 seconds. MS spectra were acquired with the Orbitrap in the range of m/z 400-1600 at a FWHM resolution of 120 000 measured at 400 m/z. AGC target was set at 4.0e5 with a 50 ms Maximum Injection Time. For internal mass calibration the 445.120025 ions were used as lock mass. The more abundant precursor ions were selected and collision-induced dissociation fragmentation was performed in the ion trap to have maximum sensitivity and yield a maximum amount of MS/MS data. Number of precursor ions was automatically defined along run in 3s windows using the “Inject Ions for All Available parallelizable time option” with a maximum injection time of 300 ms. The signal threshold for an MS/MS event was set to 5000 counts. Charge state screening was enabled to exclude precursors with 0 and 1 charge states. Dynamic exclusion was enabled with a repeat count of 1 and duration of 60 s.

### Data Processing Protocol

Relative intensity-based label-free quantification (LFQ) was processed using the MaxLFQ algorithm (Cox et al., 2014) from the freely available MaxQuant computational proteomics platform, version 1.6.3.4. The acquired raw LC Orbitrap MS data were first processed using the integrated Andromeda search engine. Spectra were searched against the Human database extracted from UniProt on the 1th of September 2020 and containing 20375 entries (reviewed). This database was supplemented with a set of 245 frequently observed contaminants. The following parameters were used for searches: (*i*) trypsin allowing cleavage before proline; (ii) two missed cleavages were allowed; (*ii*) monoisotopic precursor tolerance of 20 ppm in the first search used for recalibration, followed by 4.5 ppm for the main search and 0.5 Da for fragment ions from MS/MS ; (*iii*) cysteine carbamidomethylation (+57.02146) as a fixed modification and methionine oxidation (+15.99491) and N-terminal acetylation (+42.0106) as variable modifications; (*iv*) a maximum of five modifications per peptide allowed; and (*v*) minimum peptide length was 7 amino acids and a maximum mass of 4,600 Da. The match between runs option was enabled to transfer identifications across different LC-MS/MS replicates based on their masses and retention time within a match time window of 0.7 min and using an alignment time window of 20min. The quantification was performed using a minimum ratio count of 1 (unique + razor) and the second peptide option to allow identification of two co-fragmented co-eluting peptides with similar masses. The false discovery rate (FDR) at the peptide and protein levels were set to 1% and determined by searching a reverse database. For protein grouping, all proteins that cannot be distinguished based on their identified peptides were assembled into a single entry according to the MaxQuant rules. The statistical analysis was done with Perseus program (version 1.6.1.14) from the MaxQuant environment (https://www.maxquant.org). The LFQ normalized intensities were uploaded from the proteinGroups.txt file. First, proteins marked as contaminant, reverse hits, and “only identified by site” were discarded. Protein LFQ normalized intensities were base 2 logarithmized to obtain a normal distribution. Quantifiable proteins were defined as those detected in at least 70% of the samples in one or more condition. Missing values were replaced using data imputation by randomly selecting from a normal distribution centered on the lower edge of the intensity values that simulates signals of low abundant proteins using default parameters (a downshift of 1.8 standard deviation and a width of 0.3 of the original distribution). To determine whether a given detected protein was specifically differential a two-sample t-test were done using permutation-based FDR-controlled at 0.01 and employing 250 permutations. The p value was adjusted using a scaling factor s0 with a value of 1.

### RNA-seq bioinformatics processing

The 53 HCT116 cells (28 PTK7-Neg / 25 PTK7-Pos) were analyzed by RNAseq and computationally processed. The program Kallisto has been used to quantify the transcript reads using the transcriptome from hg19 / GRCh37 genome as reference. The alignment was performed with the program STAR on hg19 / GRCh37 reference genome. Quality Control has been made on processed data. For each cell transcriptome we computed the mitochondrial read fraction (mitoRNA) and the ribosomal read fraction (rRNA) using the Samtools function samtools coverage and the RseQC split_bam.py script, respectively. The QualiMap and STAR program outputs provided the percentage of reads aligned on exons and the percentage of uniquely mapped reads, respectively. Following cutoffs have been applied on these quality metrics: uniquely mapped reads > 70%; exonic reads > 60%; mitoRNA < 12.5%; rRNA < 2.5%.

Consequently 14 PTK7-Neg and 21 PTK7-Pos HTC116 data from 6 different NGS runs were kept for the Differential Expression Analysis (DEA).

### Normalization and Differential Expression Analysis

From the raw count matrix, the transcripts with at least 7 non-zero samples and a mean of 3 reads by sample have been selected for the downstream analyses. We normalized the resulting 15323 gene expression matrix and managed the batch difference using the R package DESeq2, available on Bioconductor. The DEA of DESeq2 identified 1284 differentially expressed genes (DEG) (8.4%) in PTK7-Pos versus PTK7-Neg cells (1073 up-regulated and 211 down-regulated genes in PTK7-Pos) (adjusted p-value < 0.01; absolute fold change > 2).

Hierarchical clustering has been achieved on the resulting signature on rows (genes) and columns (cells) (method: Ward; row distance: Pearson correlation; column distance: Euclidean). The columns’ tree clearly separates the PTK7-Pos and PTK7-Neg cells. We decided to cut the genes’ tree in 3 clusters.

### CTC *in silico* database

To assess whether these results have any significance in CRC, we mined public CTC datasets and retrieved eight datasets containing PBMCs, CTCs as well as primary tumors and metastases (detailed in **Table S11**). We limited our analysis to three robust datasets that we have validated and containing transcriptomic profiles of blood cells and CTCs.

We compared RNA expression between cell types/presentation with violin plots and searched for gene enrichment with functional signatures.

### Autophagy induction assay

HCT116 grown for 48 hours in ULA conditions were used to measure autophagy flux. Autophagy was detected using the CYTO-ID® Autophagy Detection Kit (Enzo) according to the manufacturer’s instructions. In brief, we treated HCT116 grown under ULA conditions with rapamycin to induce autophagy or with DMSO as vehicle. Cells were harvested and incubated with the CYTO-ID® probe, which selectively accumulates in autophagic vacuoles, at 37°C for 30 minutes. They were then stained with an anti-PTK7-PeCy7 (Mylteniy) mAb and Aqua LiveDead (ThermoScientific). The analysis was performed with a LSRII® (BD). The autophagy induction index was calculated by dividing the mean fluorescence of the CYTO-ID® probe of cells exposed to rapamycin by the mean fluorescence of the CYTO-ID® probe of cells exposed to vehicle medium.

### In vitro functional assay of PTK7 endocytosis

HCT116 cells were culture in ULA condition for 48h. Consecutively cells were treated for 4h either with chlorpromazine (10µM) (Ref# C8138, Merck®) or methyl-β-cyclodextrin (50µM) (Ref# C4555, Merck®) or solvent (DMSO 1/200). Then, cells were harvested and centrifuged. Supernatants from each treatment were stored at -20°C for further assays on soluble form of PTK7 (sPTK7). Pellet fractions of cells were resuspended at 1x10E6 cells/ml in PBS 1x for flow cytometry analysis.

### In vitro functional assay of PTK7 shedding

To evaluate the role of metalloproteases in PTK7 shedding, HCT116 cells were cultured under ULA conditions for 48 hours in the presence of inhibitors (Marimastat Ref# *2631,* Bio-Techne®, TMI-1 Ref# HY-101448, MedChemExpress®, NSC405020 Ref# HY-15827, MedChemExpress®, or GW280264X Ref# 7030/10, Bio-Techne®) at concentrations of 2 µM, 0.2 µM, or 0.02 µM, or with solvent (DMSO 1/500). Cells were then washed and resuspended at 1 × 10^6 cells/ml in 1× PBS for subsequent flow cytometry analysis.

To highlight the role of MMPs in PTK7 shedding, HCT116 cells were grown under ULA conditions for 48 hours, then incubated for 2 hours with PMA (Ref #HY-18739, MedChemExpress®), GW280264X, PMA + GW280264X, or solvent. Cells were then resuspended at 1 × 10^6 cells/ml in 1× PBS and processed for flow cytometry analysis.

### RNA extraction

Total RNA was isolated using the RNeasy Micro or Mini kit (Qiagen) according to the manufacturer’s instructions and treated with Dnase1. Total RNA was quantified on NanoDROP spectrophotometer ND-1000 (Thermo Fisher Scientific).

For fluidics experiments, Easy Dead Cell Removal Annexin V kit (Stemcell Technologies) was used to remove dead cells before extraction.

### Quantitative RT-PCR

Reverse transcription was performed with SuperScript™ II Reverse Transcriptase (ThermoFisherScientific). The reverse transcribed product was used to run real time PCR reactions using SYBR Green mastermix (ThermoFischer Scientific) on a Biorad CFX96^TM^ real-time cycler. Primers obtained from Life Technologies/ThermoFischer Scientific are: *PTK7 Forward* 5’- CAGTTCCTGAGGATTT CCAAGAG-3’ and *PTK7 Reverse* 5’-TGCATAGGGCCACCTTC-3’. Results were normalized to 3 reference genes: Beta2 microglobulin (*B2M Forward* 5’-GTCTTTCAGCAAGGACTGGTC-3’ and B2M Reverse 5’-CAAATGCGGCATCTTCAA ACC-3’), beta-actin (*ACTB Forward* 5’-CCACCGCGAGAAGATGA-3’ and *ACTB Reverse* 5’-CCAGAGGCGTACAGGGATAG-3’) and HPRT (*HPRT Forward* 5’-TATGGCGACC CGCAGCCCT-3’ and *HPRT Reverse* 5’-CATCTCGAGCAAGACGTTCAG-3’).

### ELISA

Ninety-six well plates (MAXISORP, NUNC) were coated with the capture antibody anti-PTK7 (clone 31G9) at 50µg/mL in PBS1X and incubated overnight at 4°C. Wells were blocked with 150µL of 1% BSA in PBS1X at room temperature for 2h. A range of soluble PTK7 (sPTK7) dilutions (from 0,49ng/mL to 1µg/mL) and samples of interest were incubated at room temperature for 2h. Biotinylated anti-PTK7 antibody (clone 10H12) diluted at 2µg/mL in PBS-1%BSA was added to each well and incubated at room temperature for 2h. Streptavidin-HRP (R&D systems, 1:200 in PBS-BSA1%) was incubated for 30min at room temperature, and finally 100µL of TMB substrate (R&D systems) was added and incubated for 5min at room temperature. Between each step, wells were washed 3 times with PBS1X-0,05%Tween and 2 times with PBS1X. To stop the color development a solution of H2SO4 2N was added and absorbance was measured at 450nm with a CLARIOstar Plus microplate reader (BMG Labtech).

### Soluble receptors Proteome profiler

Soluble receptors were analyzed using the Human Soluble Receptor Proteome Profiler Array (Non-Hematopoietic Panel, R&D Systems) according to the manufacturer’s instructions. Briefly, HCT116 cells were cultured either in 2D monolayers or as 3D spheroids and treated or not with chlorpromazine (CPMZ or TMI-1). Cell culture supernatants were collected, centrifuged to remove debris, and 500µL were incubated overnight at 4 °C with array membranes. After washing, membranes were incubated with biotinylated detection antibodies and streptavidin–HRP. Signals were detected by chemiluminescence and quantified by densitometric analysis. Data are presented as relative protein expression levels.

### Intra-venous injections models

Two types of approaches have been developed to study metastasis occurrence experimentally, which are described below.

In the first approach, HCT116-GFP-Luc cells were cultured for 24 hours under ULA conditions, then harvested, stained with PTK7 antibody (130-112-679, Myltenyi Biotec) and sorted according to their PTK7 protein expression status using the FACSAria™ III sorter (BD Biosciences).

Two groups of 6-week-old NSG mice (n=5 mice per group) were intravenously injected with 2.10^4^ sorted cells (each PTK7- or PTK7+) suspended in 100 µL PBS 1X.

In the second approach, HCT116-GFP-Luc cells were pretreated with either TMI-1 (10µM) or vehicle (DMSO 0.5%) for 24 hours. Harvested cells were counted and adjusted to a concentration of 2.10^5^ cells/mL in PBS1X. 100 µL of the pretreated cell suspension was injected intravenously into 6-week-old NSG mice (treated with TMI-1, 100 mg/Kg or vehicle 0.9% NaCl, 0.5% methylcellulose,2% Tween 80 for 24h and 1h prior injection).

In procedures that included animal follow-up, each mouse was monitored for cell diffusion efficiency by bioluminescence after intravenous injection with PhotonIMAGER™ (BIOSPACELab). Mice were examined twice weekly for 4 weeks for the appearance of metastases using bioluminescence. A longitudinal comparison of the bioluminescence signal was performed in each group of mice by comparing each time point with the background signal (first measurements within 14 days after injection, measured at ∼300 cpm). For each time point, a lateral comparison of the bioluminescent signal between the groups of mice (PTK7+ vs. PTK7-) was performed.

After successive detection of strong bioluminescent signals in the control group, mice were scarified (≈ day 30 post-injection) and organs were harvested and examined for metastatic foci by bioluminescence and analyzed using M3 Vision™ software (BIOSPACELab).

### Statistical analysis

All graphs and statistical analyses were done using the GraphPad Prism™, R studio, String™ or Excel™ software. All data are presented as mean of multiple experiments (+/- standard deviation). Statistical tests used for determination of *p*-values are specified in corresponding figure legends. For all analyses, only *p*-values <0.05 were considered as statistically significant, the key for asterisk placeholders for *p*-values in the figures are: \*\*\**p*<0.0005, ** *p* <0.005, * *p* <0.05, ns= non-significant.

## Supporting information

Supplemental Figure 3bis

## Acknowledgements

**General**

First of all, we would like to dedicate this manuscript to our mentor, Dr D Birnbaum^†^, and we are all sad that he never got the chance to see it published. We would like to also thank all the patients who contributed to the study; the Direction de la Recherche Clinique for the management of the CTC colon cohort and Dr JL Raoul, the first investigator of this cohort; Dr S. Garcia (AP-HM) and Dr M. Gilabert^†^ (IPC) for their support and expertise; The Centre de Ressources Biologiques (CRB, A Malzac) of the Paoli Calmettes institute and the Biotheque of AP-HM (E Douguy). F. Mallet and M. Richaud of the flow cytometry platform for their outstanding expertise; M. Rodrigues of the microscopy and scientific imaging platform; the histopathology platform (ICEP), the CRCM animal facility and Target; Proteomics analyses were done using the mass spectrometry facility of Marseille Proteomics (marseille-proteomique.univ-amu.fr) supported by the Institut Paoli-Calmettes and the Centre de Recherche en Cancérologie de Marseille and by IBISA (Infrastructures en Biologie Santé et Agronomie), Plateforme Technologique Aix-Marseille University, the Canceropôle PACA, the Provence-Alpes-Côte d’Azur Région and the Fonds Européen de Développement Régional (FEDER). This manuscript was edited at *Life Science Editors*.

^†^ *in memoriam*.

## Authors contribution

Conceptualization: FB, DB, JPB, EM. Methodology: JPB, EM. Clinical assays design and ethical authorization for studies involving patients: CG, DJB, FB, EM, CCM, BC, BL, AG. Patients’ enrollment and study completion in accordance to the Good Clinical Practice: DJB, CdC, BC, BL, CG, AG, FB, EM. Collect, analyze and/or interpret the data: OC, AA, AL, CA, PF, CD, LG, QdC, CG, LM, AE, SA, LC, BdR, LB, GL, ED, MP, AC, SM, EM. Formal analysis: OC, AA, AL, CA, PF, CD, LG, QdC, CG, LM, AE, SA, LC, BdR, LB, GL, ED, MP, AC, SM, EM. Validation: OC, AA, AL, CA, PF, CD, LG, QdC, CG, LM, AE, SA, LC, BdR, LB, GL, ED, MP, AC, SM, EM; Data curation: OC, AA, AL, CA, PF, EM. Funding acquisition: DB, FB, DJB, JPB, EM. Project administration: DB, FB, DJB, JPB, EM; Supervision: JPB, EM. Writing – original draft: OC, AA, AL, CA, EM. Writing – finalize, review & editing: DB, FB, JPB, EM. All authors have read and approved the final version of the manuscript and agree to be accountable for all aspects of the work.

## Data and materials availability

The transcriptomic data are publicly available and details about repository and reference are listed in **Table S1**.

The mass spectrometry proteomics data have been deposited to the ProteomeXchange Consortium via the PRIDE partner repository under the dataset identifiers PXDxxxxx, with the following reviewer account details: Username: / Password: xxxxxx.

All the other data generated in this study are available within the article, its supplementary data files or on request.

## Ethics approval and consent to participate

In the present study, samples from the prospective trials CTC-Colon and B-Org were used, which are registered under the https://ClinicalTrials.gov identifier NCT03256084 and NCT05384184, and approved by the French National Agency for the Safety of Medicines and Health Products, a national ethics committee (CPP Sud-Méditerranée, under the study ID number ID RCB: 2016-A00621-50 and 2019-A00710-57), the Paoli-Calmettes Institute Institutional Review Board (Comité d’Orientation Stratégique or COS, under the study ID number CTC-Colon-IPC 2015-020) and the AP-HM (2019-A00710-57). The protocols are available at https://clinicaltrials.gov/study/NCT03256084 and https://clinicaltrials.gov/study/NCT05384184. The studies were conducted in accordance with the Declaration of Helsinki. All patients gave informed consent for inclusion, biopsy and blood sampling, and molecular analysis of these samples. Patient privacy was protected at all times during the study. Informed consent was obtained from all persons involved in the study.

## Supplementary Materials

**Table S1:** The 11 colon cancer data sets used in this study.

**Table S2:** Clinicopathological characteristics and correlations with the PTK7 High/Low-based classification.

**Table S3.** Univariate and multivariate analyzes for OS.

**Table S4.** List of 316 genes that are differentially expressed between the PTK7^high^ and PTK7^low^ classes. Supervised analysis COAD TCGA 367 up vs. 92 no-up PTK7 (moderated t-test, threshold: p<5%, q<25% & |FC|>2x)

**Table S5.** Gene ontologies associated with the 316 genes that are differentially expressed between the PTK7^high^ and PTK7^low^ classes.

**Table S6.** Clinicopathological characteristics Clinical data of the CTC-colon cohort.

**Table S7.** Surfaceome proteomics analysis of HCT116 cells in non-injected conditions, subjected to a fluidic circulation system, and following recovery and re-adhesion.

**Table S8.** List of 1284 genes that are differentially expressed between the PTK7^+^ and PTK7^-^ cells. Supervised analysis 21 PTK7^+^ vs. 14 PTK7^-^ single cells (moderated t-test, threshold: p<5%, q<25% & |FC|>2x)

**Table S9.** Gene ontologies associated with the 1284 genes that are differentially expressed between the PTK7^+^ (Orange) and PTK7^-^ (Blue) phenotype.

**Table S10.** Antibodies and labelling reagents

**Table S11.** Details of the 8 public data sets used to construct the “CTC database”.

**Sup Figure 1: PTK7*^high^* expression in CRC is associated with poorer survival of patients with the mesenchymal subtype of CRC.**

**A.** Localization of PTK7 at the base of the crypt in normal colon tissue measured by IHC.

**B.** Boxplots showing *PTK7* mRNA expression levels (log_2_ compared to normal colon) in primary CRC by CMS subtypes. CMS1 (n = 389, yellow), CMS2 (n=640, blue), CMS3 (n=343, pink), CMS4 (n=604, green). *PTK7* expression was compared between groups using ANOVA (multiple comparisons). **C.** Kaplan-Meier OS curves of patients with primary CRC according to PTK7-based classification by CMS subtypes. The p-values are for the log-rank tests in the global population and for each CMS subtypes. **D**. EMT and tumor cells budding metagenes enrichment (GSEA EMT hallmark metagene V62, EMT/CHL model, EMT score, tumor cell budding signature, and C1 to C4 EMT states) in *PTK7^Low^* and *PTK7^High^*tumor groups (transcriptomic level). **E**. Expression of PTK7 by IHC in tumor invasion clusters, tumor buds and small tumor glands. **F.** Expression of PTK7 (mean fluorescence intensity) in HCT116 and SW480 cell lines exposed to StemX Vivo®, an EMT inducing agent. **G**. Immunofluorescence PTK7 labelling following a scratch assay.

**Sup Figure 2: PTK7 expression on primary tumor cells does not presuppose PTK7 expression on CTCs.**

**A.** Representative images of immunofluorescence staining of CTC clusters isolated from the blood of 2 CRC patients. CTCs are recognized by the expression of the epithelial markers EPCAM and/or pan-CK and/or VIMENTIN (VIM) and by the absence of CD45. The expression of PTK7 (here, in red) was assessed on these cells. **B.** An example of PTK7 staining score 3 by IHC in a primary CRC. **C.** Number of CTCs analyzed according to the PTK7- or PTK7+ phenotype in patients with a PTK7 staining score of 3 in their primary CRC tissues (n = 8 patients). **D.** Number of CTCs analyzed according to the PTK7- or PTK7+ phenotype in patients whose CRC had progressed to a metastatic stage within 2 years of enrollment in our study (n = 7 patients). **E**. Summary of the PTK7 status (positive or negative) by IHC in mice primary tumors and the corresponding metastatic sites, for each of the 6 mice. **F**. Gating strategy for the detection of CTCs in mouse blood samples. An initial gate was used to select large cells and avoid cellular debris (P1). These events were then plotted against the mouse histone H2 and a cell death marker (Livedead™). The double negative gate P2 was used to exclude H2+ mouse cells and dead or dying cells (P2). Finally, events selected in P2 were plotted for GFP + anti-human EPCAM-A488 and PTK7 to select epithelial cells of human origin and determine the corresponding phenotype for PTK7.

**Sup Figure 3: PTK7 ON^tumor^/OFF^CTC^/ON^metastasis^ switch is influenced by the mechanical constraints experienced by CTCs during fluidic stress.**

**A.** Surfaceome proteomic profiling of HCT116 CRC cells was performed under three conditions: non-injected adherent cells, cells exposed to a 4-hour fluidic circulation system (circulating cells), and cells allowed to re-adhere after circulation (re-adherent cells). Gene Set Enrichment Analysis (GSEA) of KEGG pathways performed using differential surfaceome proteomic data obtained from the comparison between non-injected adherent cells versus circulating cells, and circulating cells versus re-adherent cells. Each dot represents a significantly enriched pathway. The x-axis indicates the normalized enrichment score (NES). **B.** Gene-pathway interaction network generated from significantly enriched KEGG pathways identified in the comparison between non-injected adherent and circulating cells. Protein node color indicates the relative fold change between the compared conditions. **C.** Gene-pathway interaction network of KEGG pathways altered between circulating cells versus re-adherent cells**. D**. Representative images of immune-fluorescent staining of PTK7 (in orange) in HCT116 and SW480 cell lines. Scale: 10µm. **E**. Schematic representation of the different culture conditions used in the study (top) and percentage of SW480 cells expressing PTK7 when exposed to the indicated adhesion conditions (bottom). A two-way ANOVA with a Bonferroni post-tests was used to compare the conditions between them. **F.** Representative flow cytometry histograms showing PTK7 expression on SW480 cells in the control condition (purple) or after 24 and 48 hours of culture in ULA + Ca^2+^ medium (green) or in flow (orange) conditions. **G**. Representative images of immunofluorescence staining of PTK7 (green), EPCAM (red) and DAPI (blue) on HCT116 cells grown for 48 hours in indicated conditions. **H**. Bar plots of PTK7 and EPCAM fluorescence intensity in viable HCT116 cells grown in ULA - Ca2+ or after flow exposition. Fluorescence intensity is expressed as a percentage compared to the control condition, *ie*. to cell grown in 2D + Ca2+ (n = 6 experiments). **I**. A representative image of immunofluorescence staining of a HCT116 spheroid growing in the upper part of the Boyden chamber (DAPI (blue) and PTK7 (green)). **J**. Representative flow cytometry histograms showing PTK7 expression on HCT116 grown in 2D, representing the control condition, after 48 hours of flow exposure and after 48 hours of culture in adherent condition following flow exposure.

**Sup Figure 4: PTK7- cells activate the YAP1 signaling pathway.**

**A.** Workflow recapitulating the key steps to isolate PTK7+ and PTK7- cells before performing the single-cell RNAseq analysis. **B**. Enrichment of the hallmark cancer gene expression signatures (top) and stress response signatures (bottom) in PTK7+ and PTK7- cells. **C**. Expressions of PTK7 (green) and CTGF (red) in HCT116 cells before and after fluidic experiments, nuclei were stained with DAPI (Blue) (representative images of 3 independent experiments).

**Sup Figure 5: The shedding of PTK7 involves the activation of MT1-MMP and ADAM17.**

**A.** Correlation between *PTK7* mRNA levels measured by RT-qPCR in HCT116 exposed to various conditions and PTK7+ expression measured by FACS. *PTK7* mRNA level WiDr cell line was added as a reference for PTK7 low expression. **B**. PTK7 protein expression in WiDr cell line compared to HCT116 cell line. Position of the control isotype for each cell lines is represented by a grey histogram. **C**. Representation of the full-length form of PTK7, with the known cleavage sites by MT1-MMP, ADAM17 and γ-secretase indicated. *Adapted from* Golubkov and Strongin, 2012 *et al.*.

**D.** Percentage of PTK7- HCT116 cells cultured in ULA +Ca^2+^ medium and treated with methyl-β-cyclodextrin or DMSO (vehicle control). **E**. Representative flow cytometry histograms of the effect of chlorpromazine on PTK7 expression (vehicle-treated cells on the left and chlorpromazine-treated cells on the right). **F**. Quantity of sPTK7 detected by ELISA in the plasma of healthy subjects, CRC patients with primary or metastatic disease. **G**. Correlation between sPTK7 (ng/ml) in the plasma and the number of CTCs/ml detected in the corresponding CRC patients. **H**. Quantity of sPTK7, sMT1-MMP, sADAM17 detected by ELISA in the plasma of CRC patients. **I**. Percentage of PTK7- cells observed after co treatment with decreasing concentration of MMPs inhibitors. **J**. Representative flow cytometry histograms after HCT116 cells treated with ADAM17 inhibitor (GW280264X), with or without activation with PMA. **K**. Results of the soluble receptor proteome profiler® array on the supernatant of HCT116 cells grown in ULA- Ca2+ and in 2D. Results are expressed as the fold change between these two conditions.

Results are expressed as the mean of at least 3 independent experiments ± SD. Correlations were established with a Pearson’s test. P-values were determined using a two-way ANOVA (*p <0.05).

**Sup Figure 6: PTK7- cells exhibit higher metastatic efficiency.**

**A.** Number of bioluminescent foci observed per organ in the *in vivo* experiments. **B**. Number of organs with bioluminescent foci in the mice injected with PTK7+ and PTK7-HCT116 cells. **C.** Number of organs with bioluminescent foci in the mice treated with vehicle or TM1-1 and injected with HCT116 cells. Data are expressed as mean +/- SD. P-values were determined using a one-tailed non-parametric t-test (*p <0.05).

