## Supplementary figures and images for "Metalloprotease-driven remodeling of PTK7 and the cell surfaceome promotes metastatic fitness of circulating colorectal tumor cells"

### Supplemental Figure 1

#### Figure S1

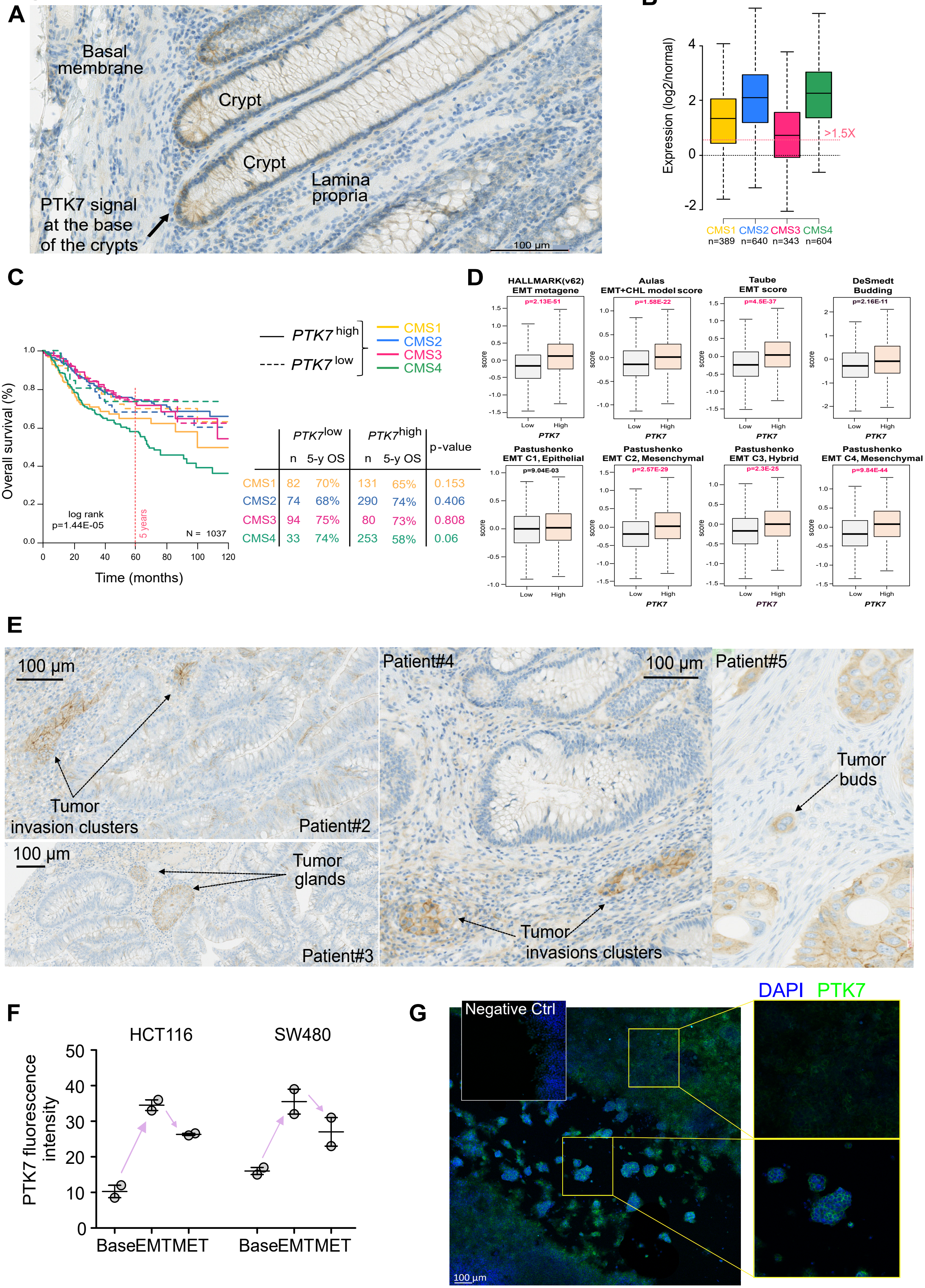


| | $PTK7^{low}$ n | $PTK7^{low}$ 5-y OS | $PTK7^{high}$ n | $PTK7^{high}$ 5-y OS | p-value |
|---|---|---|---|---|---|
| CMS1 | 82 | 70% | 131 | 65% | 0.153 |
| CMS2 | 74 | 68% | 290 | 74% | 0.406 |
| CMS3 | 94 | 75% | 80 | 73% | 0.808 |
| CMS4 | 33 | 74% | 253 | 58% | 0.06 |

### Supplemental Figure 3

#### Supplementary Figure 3 (Part 1)

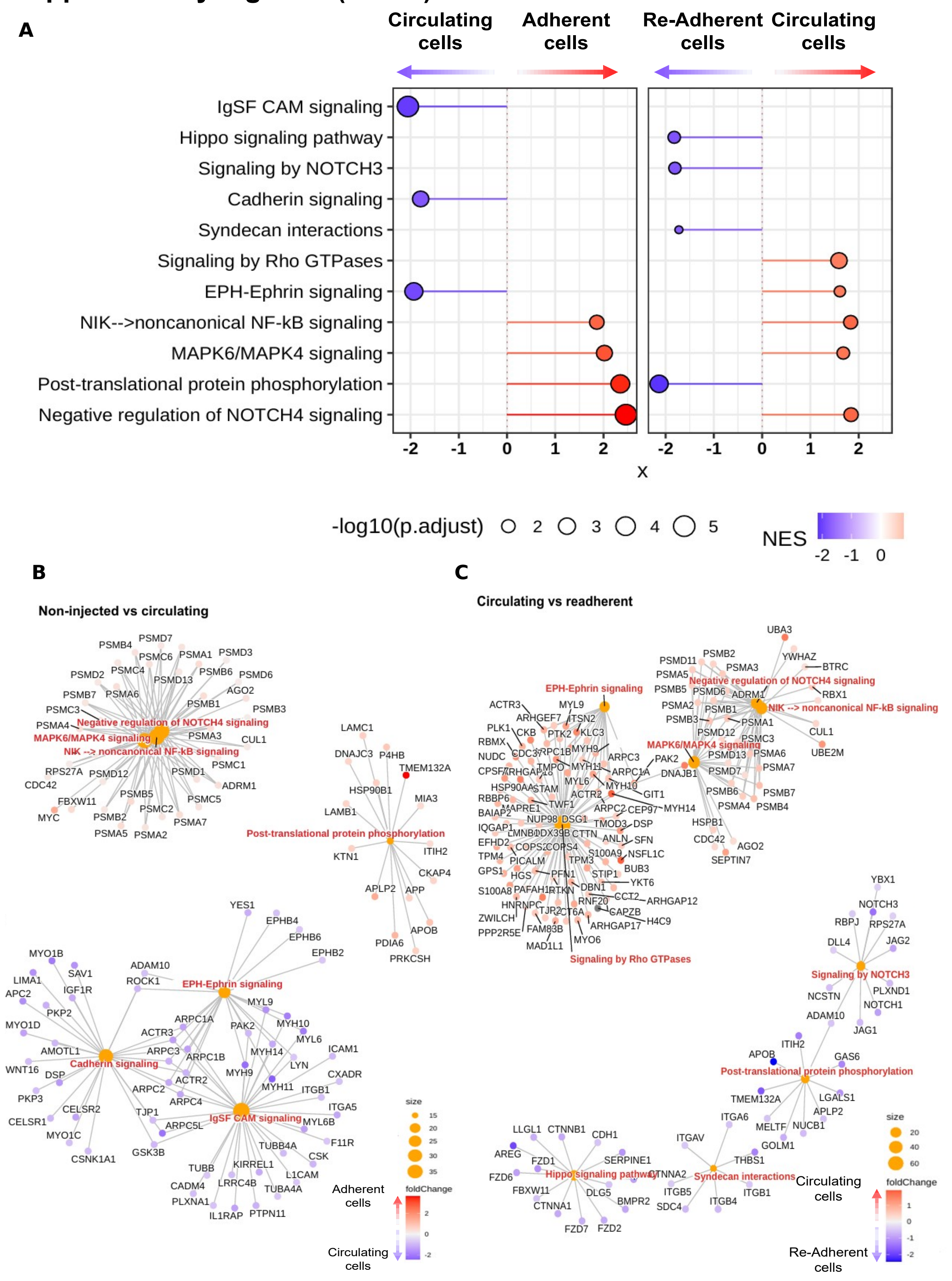

### Supplemental Figure 3bis

#### Supplementary Figure 3 (part 2)

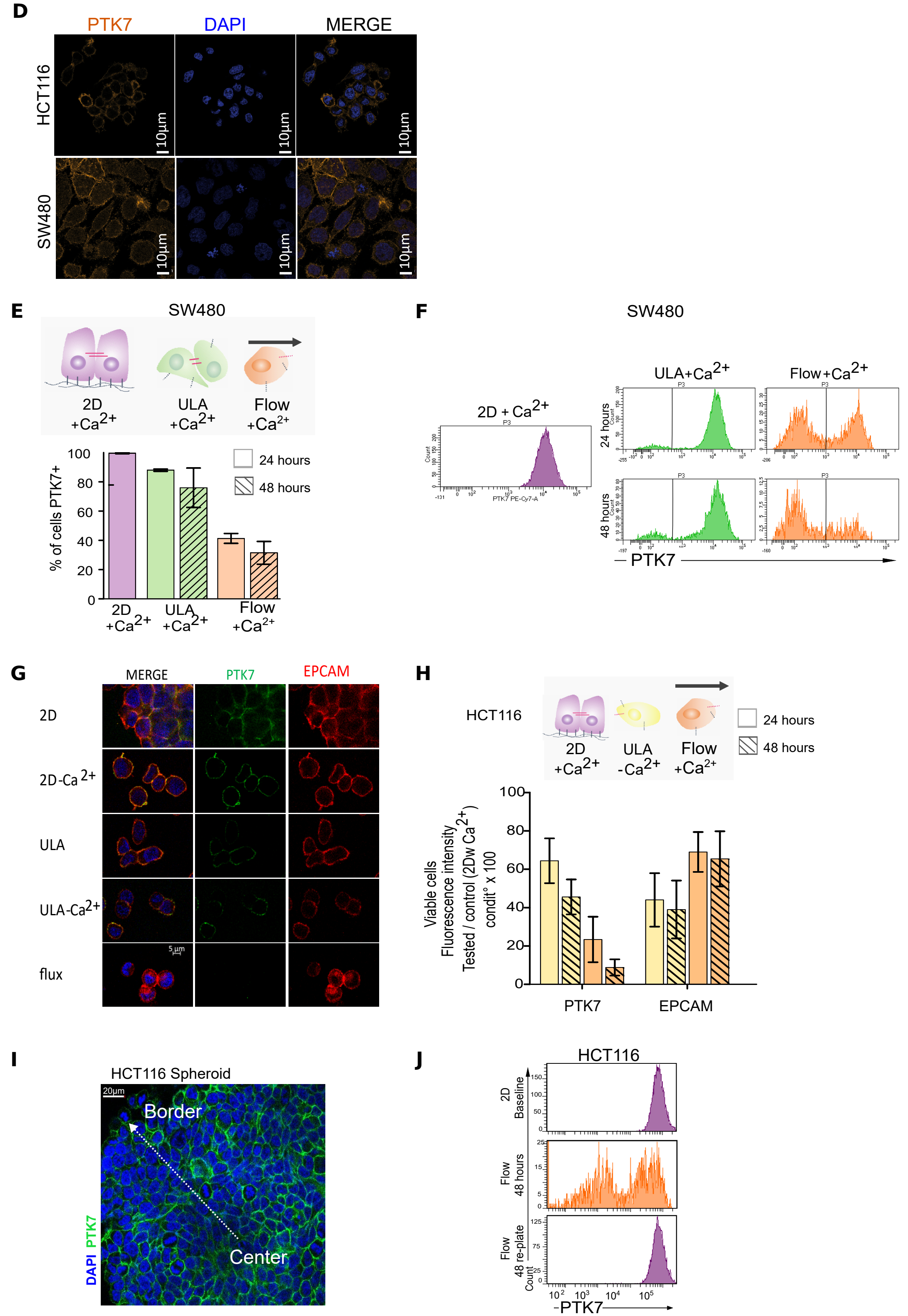

### Supplemental Figure 4

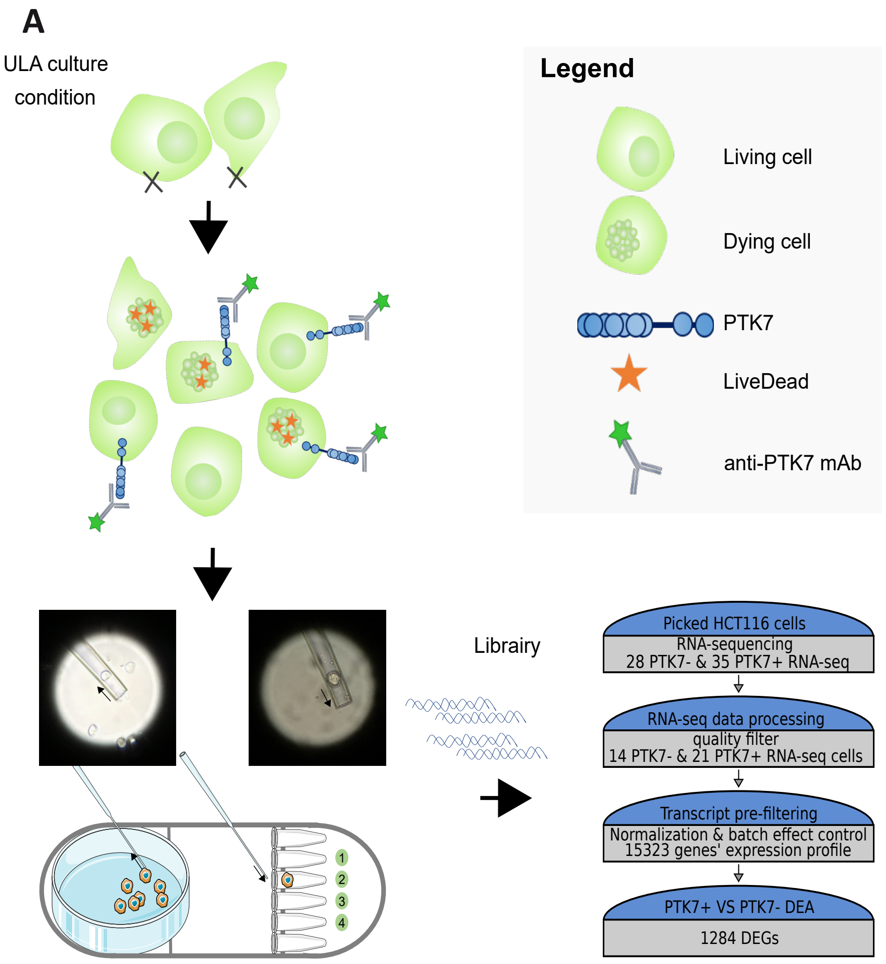

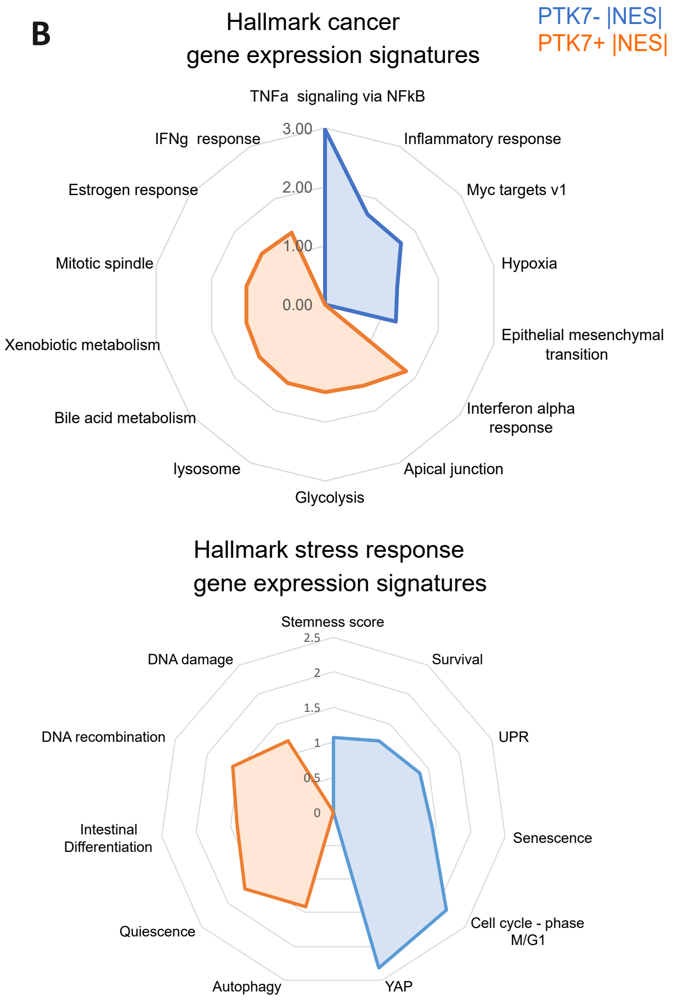

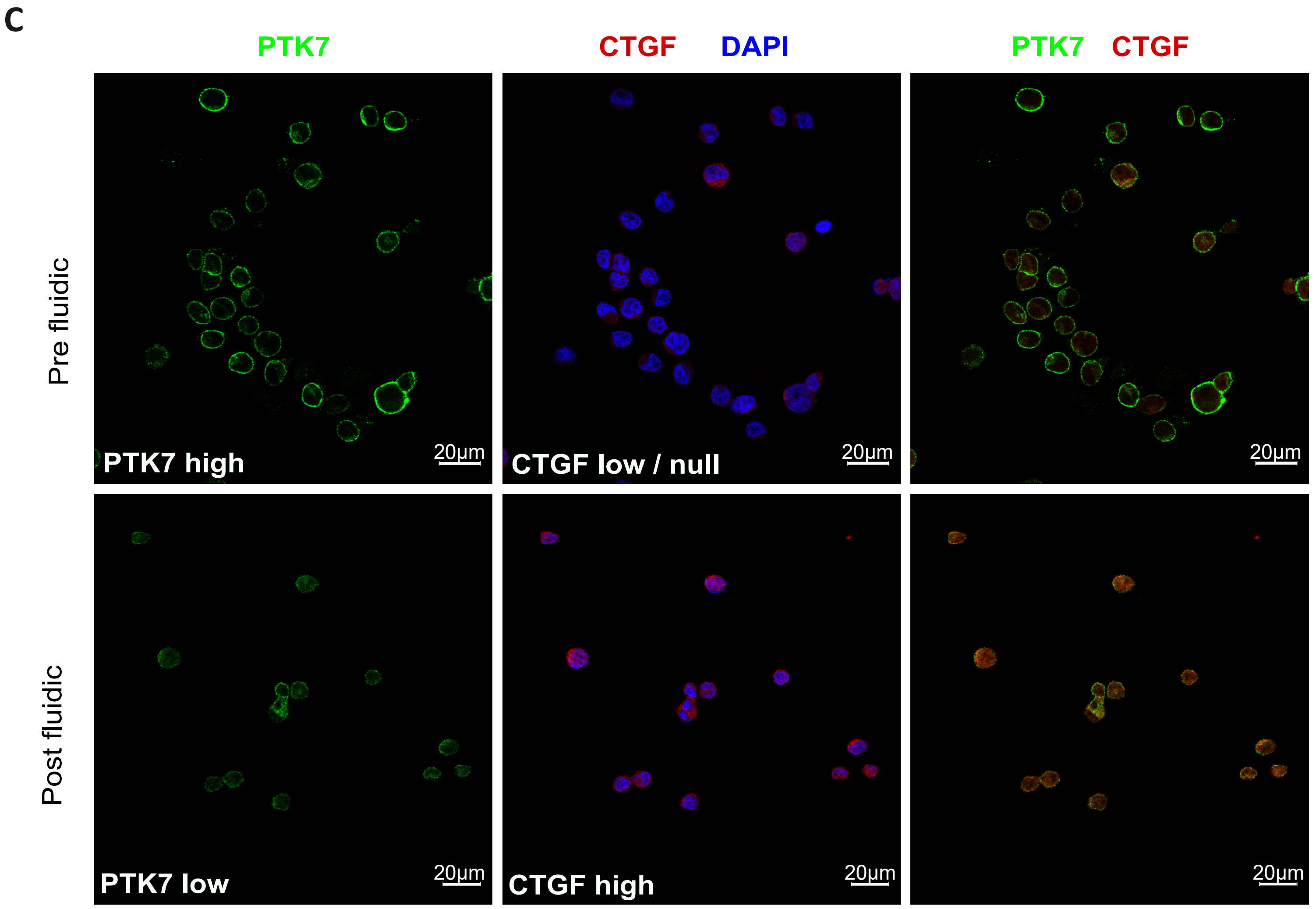

### Supplemental Figure 5

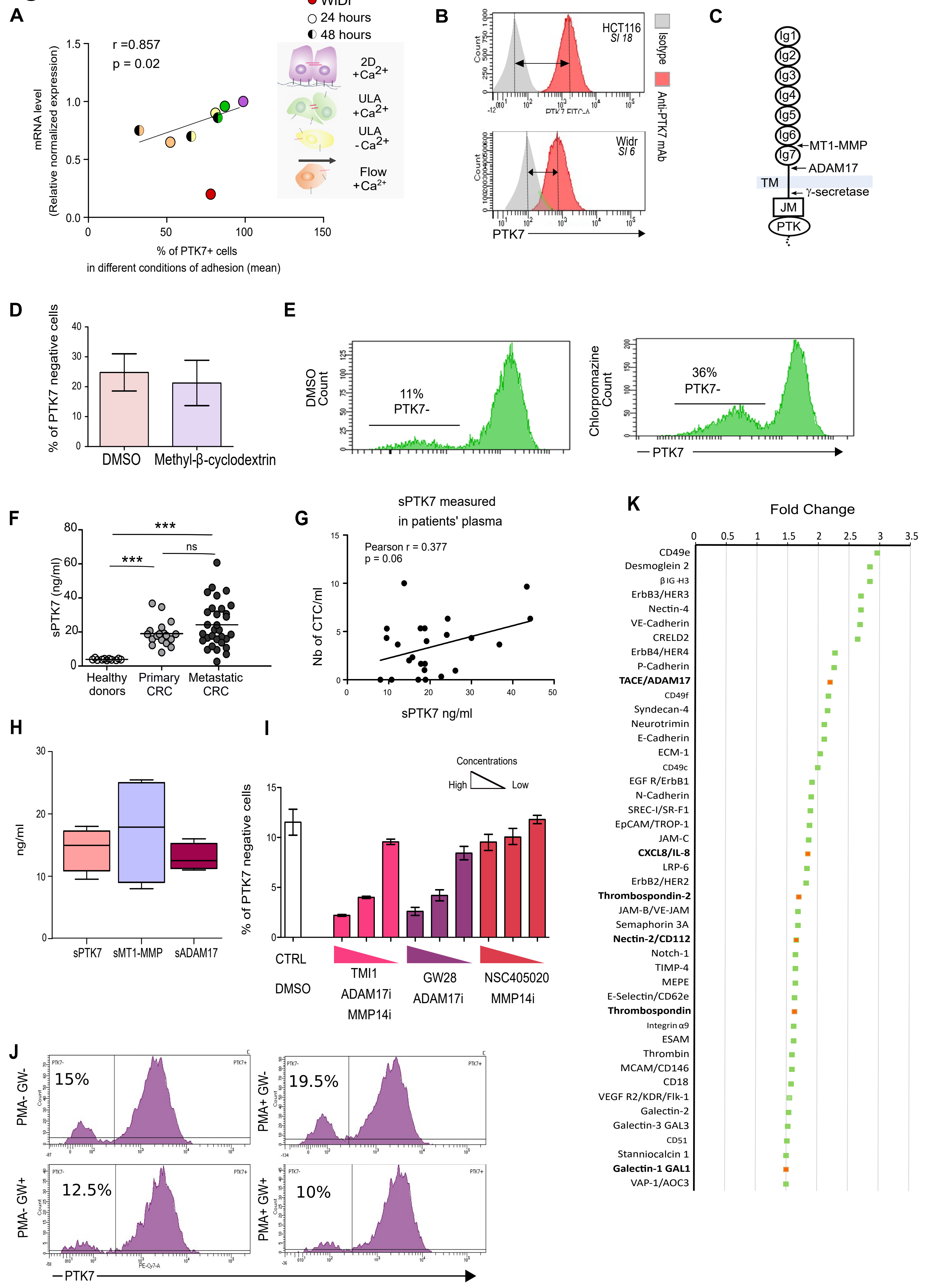
