## Supplemental Figure 2 for "Metalloprotease-driven remodeling of PTK7 and the cell surfaceome promotes metastatic fitness of circulating colorectal tumor cells"

**Figure S2**

**A**

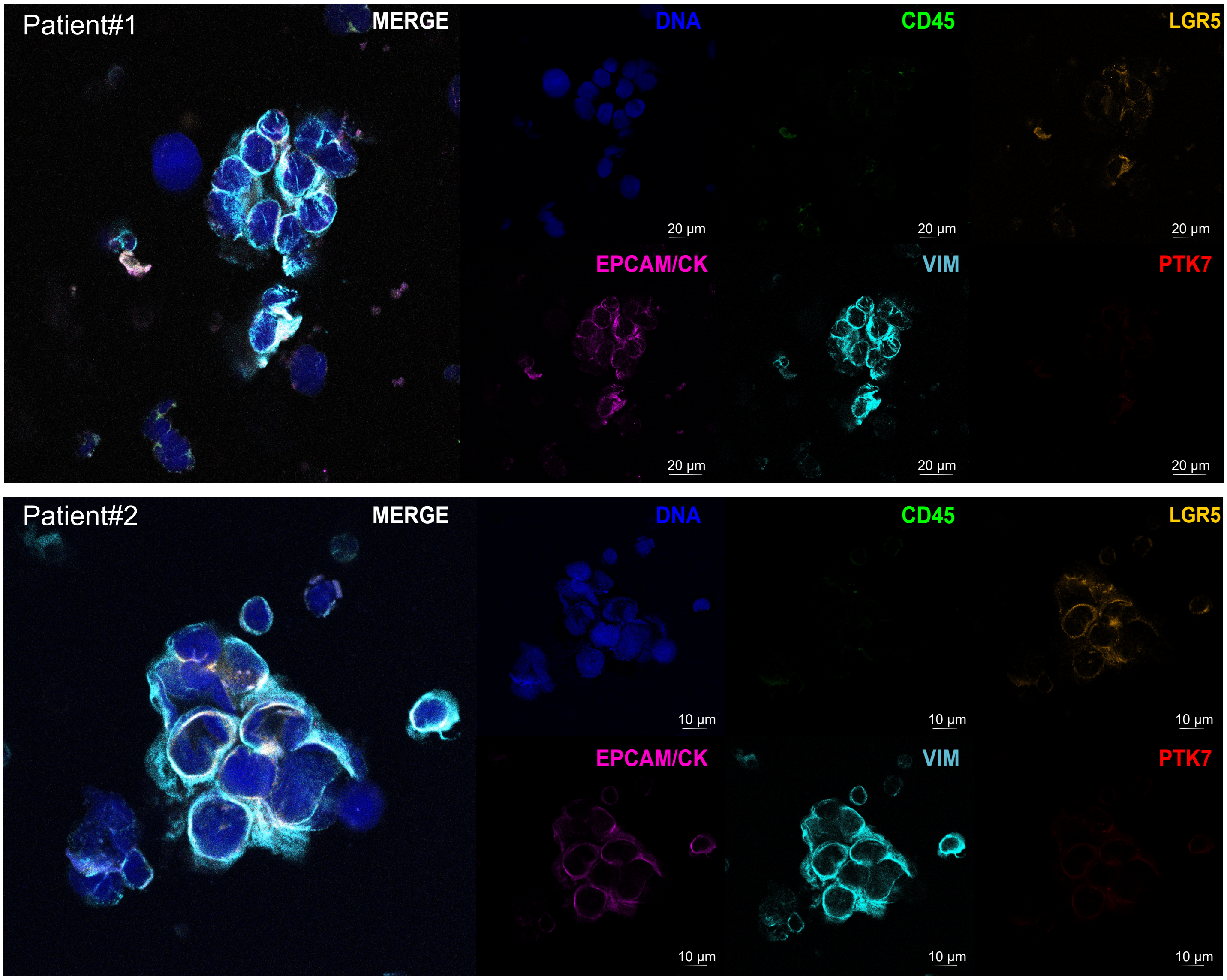


**B**

Exemple of IHC score 3 tumor

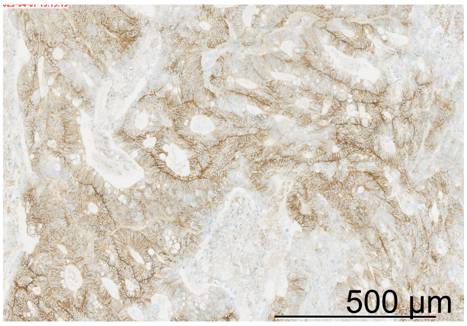


**C**

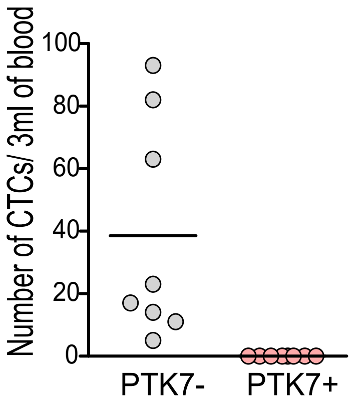


**D**

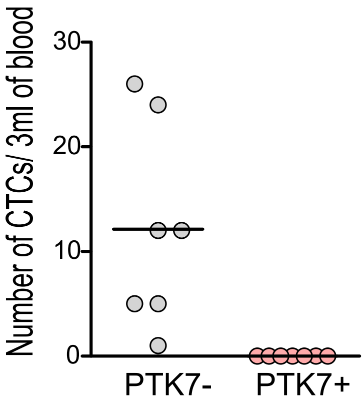


**E**

| Mouse ID | PTK7 status at primary site (IHC) | Metastatic sites (Bioluminescence) | PTK7 positivity of the metastatic sites tested (IHC) |
|---|---|---|---|
| 1 | Positive | Lymph nodes, lung, pancreas, uterine horn | 4/4 |
| 2 | Positive | Pancreas, uterine horn, liver | 2/2 |
| 3 | Positive | Lymph node, lung, pancreas, uterine horn, liver, spleen | 3/3 |
| 4 | Positive | Lymp node, lung, pancreas | 1/1 |
| 5 | Positive | Pancreas, ovary, liver, kidney | 3/3 |
| 6 | Positive | Lung, pancreas, ovary, liver | 1/1 |

**F**

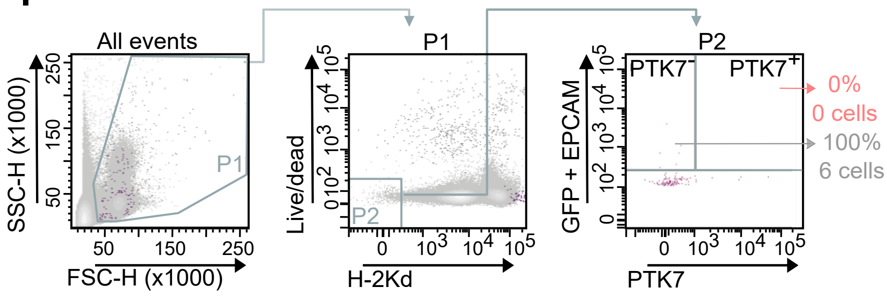
