## Supplemental Figure 6 for "Metalloprotease-driven remodeling of PTK7 and the cell surfaceome promotes metastatic fitness of circulating colorectal tumor cells"

### Figure S6

Number of bioluminescent foci per organ

**A**

| | Lungs | Liver | Kidneys | Pancreas | Ovaries | Uterin horns | Brain | Spleen | Cervical lymph nodes | Axill./brach. Lymph nodes | Mesenteric lymph nodes | Inguinal lymph nodes | Total number of bioluminescent foci |
|---|---|---|---|---|---|---|---|---|---|---|---|---|---|
| **PTK7+ (n=5)** | 9 | 8 | 2 | 2 | 1 | 0 | 0 | 0 | 2 | 0 | 3 | 2 | 29 |
| **PTK7 - (n=5)** | 20 | 10 | 5 | 1 | 2 | 0 | 0 | 1 | 5 | 2 | 2 | 5 | 53 |
| **Vehicule (n=4)** | 16 | 6 | 3 | 0 | 3 | 1 | 0 | 0 | 0 | 0 | 1 | 2 | 32 |
| **TMI-001 (n=4)** | 3 | 1 | 0 | 0 | 0 | 0 | 0 | 0 | 0 | 0 | 0 | 0 | 4 |

**B**

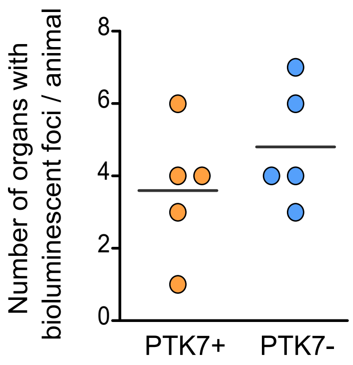


**C**

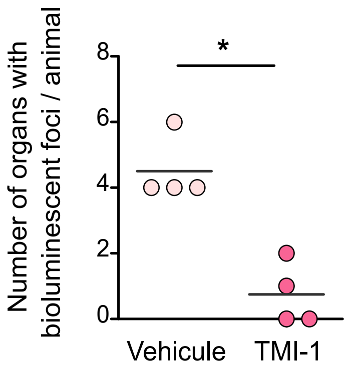
